# Dynamic integration of forward planning and heuristic preferences during multiple goal pursuit

**DOI:** 10.1101/838425

**Authors:** Florian Ott, Dimitrije Marković, Alexander Strobel, Stefan J. Kiebel

## Abstract

Selecting goals and successfully pursuing them in an uncertain and dynamic environment is an important aspect of human behaviour. In order to decide which goal to pursue at what point in time, one has to evaluate the consequences of one’s actions over future time steps by forward planning. However, when the goal is still temporally distant, detailed forward planning can be prohibitively costly. One way to select actions at minimal computational costs is to use heuristics. It is an open question how humans mix heuristics with forward planning to balance computational costs with goal reaching performance. To test a hypothesis about dynamic mixing of heuristics with forward planning, we used a novel stochastic sequential two-goal task. Comparing participants’ decisions with an optimal full planning agent, we found that at the early stages of goal-reaching sequences, in which both goals are temporally distant and planning complexity is high, on average 42% (SD = 19%) of participants’ choices deviated from the agent’s optimal choices. Only towards the end of the sequence, participant’s behaviour converged to near optimal performance. Subsequent model-based analyses showed that participants used heuristic preferences when the goal was temporally distant and switched to forward planning when the goal was close.

**Author summary:** When we pursue our goals, there is often a moment when we recognize that we did not make the progress that we hoped for. What should we do now? Persevere to achieve the original goal, or switch to another goal? Two features of real-world goal pursuit make these decisions particularly complex. First, goals can lie far into an unpredictable future and second, there are many potential goals to pursue. When potential goals are temporally distant, human decision makers cannot use an exhaustive planning strategy, rendering simpler rules of thumb more appropriate. An important question is how humans adjust the rule of thumb approach once they get closer to the goal. We addressed this question using a novel sequential two-goal task and analysed the choice data using a computational model which arbitrates between a rule of thumb and accurate planning. We found that participants’ decision making progressively improved as the goal came closer and that this improvement was most likely caused by participants starting to plan ahead.

## Introduction

Decisions of which goal to pursue at what point in time are central to everyday life [1–3]. Typically, in our dynamic environment, the outcomes of our decisions are stochastic and one cannot predict with certainty whether a preferred goal can be reached. Often, our environment also presents alternative goals that may be less preferred but can be reached with a higher probability than the preferred goal. For example, when working towards a specific dream position in a career, it may turn out after some time that the position is unlikely to be obtained, while another less preferred position can be secured. The decision to make is whether one should continue working towards the preferred position, or switch goals and secure the less preferred position. The risk when pursuing the preferred position is to lose out on both positions. This decision dilemma ‘should I risk it and go after a big reward or play it safe and gain less?’ is typical for many decisions we have to make in real life. Critically, for many such decisions, these binary choices do not emerge suddenly and unexpectedly, but the decision maker is typically confronted with such decisions after some prolonged period of time working towards enabling different options.

How would one choose one’s actions during such a prolonged goal-reaching decision making sequence? One way, if the rules of the dynamic environment and its uncertainties are known, is to use forward planning to always choose the actions which maximize the gain (see [4, 5] reviewing cognitive processes of forward planning). This would be the way one would program an optimal agent in a game or experimental task environment. This approach is often used in cognitive neuroscience to model the mechanism of how humans make decisions in temporally extended goal-reaching scenarios, (e.g. [6–9]).

However, the implicit assumption made in these decision-making models, namely that humans use detailed forward planning and compute the probabilities of reaching the goals, is difficult to justify, because of the involved computational complexity. In a stochastic environment, forward planning in artificial agents is typically achieved via sampling many possible policies (sequences of actions) which requires substantial computing power that scales exponentially with the number of future actions. In particular, when one is still temporally far from the goal, the computational burden of simulating trajectories into the future is the largest, while the usefulness of the resulting action selection is minimal: intuitively, in stochastic and sufficiently complex environments, anything may yet happen on the long way to the goal so the gain of planning ahead at high cost may be small. The importance of the balance between the benefits and its costs to better understand human decision making became a recent research focus, e.g., [10–14]. The question is how one can select actions over long stretches of time, without being exposed to the computational burden of forward planning or similar dynamic programming schemes.

One obvious way to select actions at minimal computational costs is to use heuristics that do not require forward planning towards a goal [15, 16], e.g. to always select the action towards a hard to achieve and highly rewarded goal. Clearly, this and other heuristics come with the drawback that they can be substantially suboptimal when close to the goal. For example, blindly working toward a hard to achieve goal would ignore the risk of not reaching any goal. Another solution is to use habit-like strategies to avoid computational costs [17]. However, habits are typically useful only when one encounters exactly the same situation or context repeatedly, while goal reaching in uncertain environments as presented here, often requires flexible behavioural control.

It is an open question how humans select their actions when the potentially reachable goals are still far away and forward planning is complex. We hypothesized that people use a mixture of two approaches to achieve an acceptable balance between outcome and computational costs. This mixture changes with temporal distance to the goal: when far from the goal, people use a prior goal preference to make their decision about which action to take. With this approach, one assumes that one will eventually reach the preferred goal and selects the action that, if one looked backward in time from the reached goal, is the most instrumental. When coming closer to the goal, one expects that the influence of the goal preference should be progressively superseded by computationally more expensive action selection using forward planning to optimally reach the preferred goal or, failing that one, to pursue policies to reach an alternative goal.

To test whether participants used such an approach, we employed a novel behavioural task where participants were placed in a dynamic and stochastic sequential decision task environment that emulated reaching goals over an extended time period. In miniblocks of 15 trials, participants had to make decisions to reach one or two goals, where reaching both goals was rewarded more than reaching only one. In each miniblock, it was also possible, if blindly trying to obtain the higher reward, to not reach any goal and not obtain any reward. While participants pass through the miniblock, both the remaining trials to the end of the miniblock and the complexity of forward planning decrease. This enables us to test and model whether participants switch from using heuristics to forward planning during goal-reaching. To analyse the behavioural data of 89 participants and test hypotheses, we used stochastic variational inference, which provided posterior beliefs about the goal strategy preference of each participant, among other free model parameters. We show that the heuristic goal strategy preference parameter is key to explain participants’ choices when temporally distant from the goal, and how, when progressing towards a goal, this goal strategy preference interacts with optimal forward planning to achieve near-optimal performance.

## Methods

### Participants

Eighty-nine participants took part in the experiment (58 women, mean age = 24.8, SD = 7.1). Reimbursement was a fixed amount of 8€ or class credit plus a performance-dependent bonus (mean bonus = 3.88€, SD = 13.6). The study was approved by the Institutional Review Board of the Technische Universität Dresden and conducted in accordance to ethical standards of the Declaration of Helsinki. All participants were informed about the purpose and the procedure of the study and gave written informed consent prior to the experiment. All participants had normal or corrected-to-normal vision.

### Experimental Task

The experiment included a training phase of 10 miniblocks, followed by the main experiment comprising 60 miniblocks. The 60 miniblocks in the main experiment were subdivided into three sessions of 20 miniblocks between which participants could make a self-determined pause. A miniblock consisted of *T* = 15 trials in which participants had to accept or reject presented offers to collect A-points 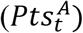 and B-points (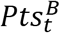, see Table 1 for a glossary of abbreviations). If participants reached the threshold of 10 points for either A- or B-point scale after 15 trials, they received a reward of 5 cents. If participants reached the threshold for both point scales, they received a reward of 10 cents. If none of the two thresholds was reached, no additional reward was provided. In total, each participant completed 150 training trials and 900 trials in the main experiment.

**Table 1.** Glossary of abbreviations

| Abbreviation | Explanation |
| --- | --- |
| A, B | Basic offers |
| Ab, aB | Mixed offers |
| $Pts_t^A$ | A-points in trial t |
| $Pts_t^B$ | B-points in trial t |
| g1 | One-goal-choice = Sequential strategy choice = Choice that maximizes point difference |
| g2 | Two-goal-choice = Parallel strategy choice = Choice that minimizes point difference |
| G1 | One-goal-success = One point scale above threshold after 15 trials |
| G2 | Two-goal-success = Both scales above threshold after 15 trials |
| Q(s,a) | Action value = Expected future reward of a choice |
| $Q_G(s,a)$ | Goal choice value = Expected future reward of a goal strategy choice |
| DEV | Differential expected value = $Q_G(s, g2) - Q_G(s, g1)$ |

Each trial started with a response phase lasting until a response was made, but not more than 3 s (Fig 1, A). The current amount of A-points and B-points was visualized by two vertical bars flanking the stimulus display. Horizontal white lines marked the threshold of 10 points. At the top of the screen, a grey timeline informed the participants about the remaining trials in the miniblock. The current offer was displayed at the bottom centre, and the two choice options were presented in the centre of the screen by the framed words ‘accept’ and ‘wait’. Participants could accept an offer by an upwards keypress and reject the offer by a downwards keypress. If participants did not respond within 3 s the trial was aborted, and a message was displayed reminding the participant to pay attention. If participants missed the response deadline more than 5 times in the whole main experiment, 50 cents were subtracted from their final payoff (mean number of timeouts = 1.34, SD = 1.7). After the response phase, feedback was displayed for 1.5 s. Response feedback included a change in colour of the frame around the selected response from white to green. Additionally, the gain or loss of points was visualized by colouring the respective area on the bar either green or red. After 15 trials, feedback for the miniblock was displayed for 4 s informing the participants whether they won 5, 10 or 0 cents. Code for experimental control and stimulus presentation was custom written in Matlab (MathWorks) with extensions from the Psychophysics toolbox [18].

**Fig 1.**
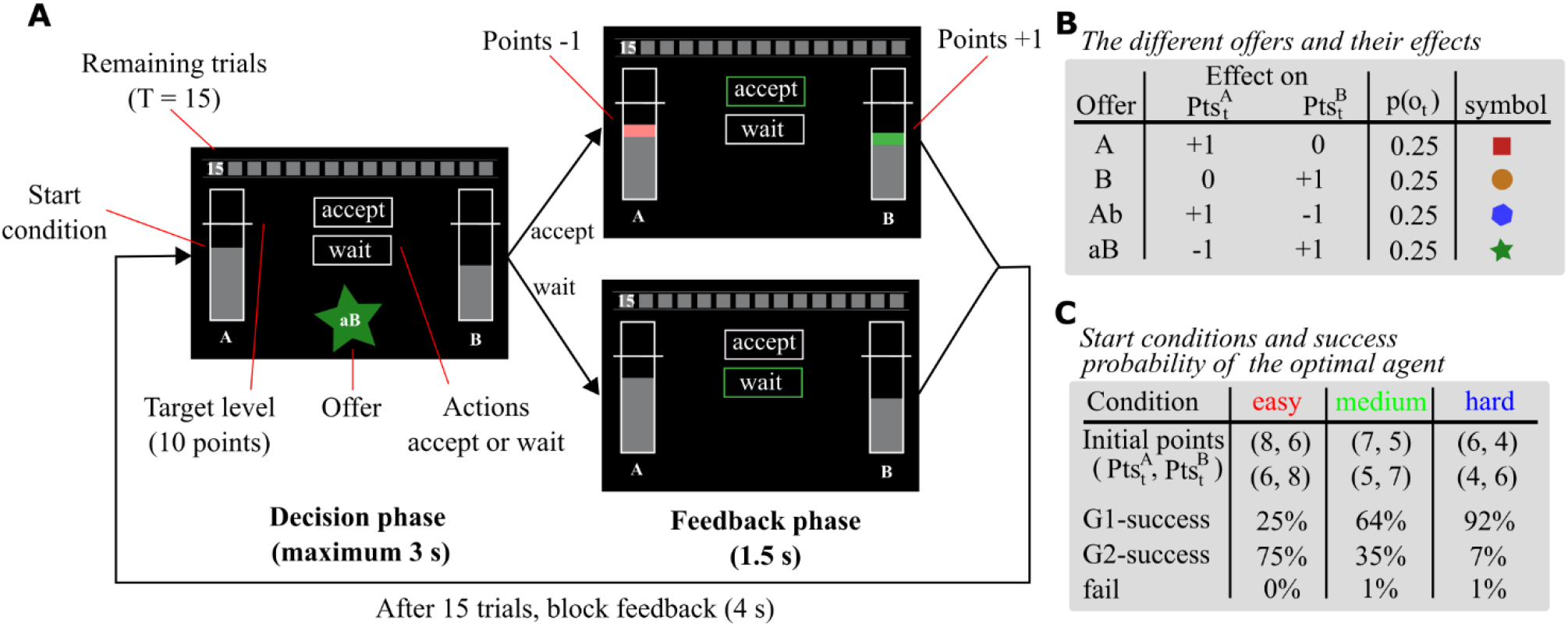
Experimental task. **(A)** Depiction of trial timeline and stimulus features. Participants performed miniblocks of 15 trials in which they collected points to reach either one or two goals, rewarding them with additional 5 or 10 Cents. Each trial started with a decision phase (maximum 3s) in which participants had to accept or reject a presented offer. Depending on the offer, accepting increased or decreased A- and B-points. The current amount of points was displayed by two grey bars flanking the stimulus screen. In the feedback phase (1.5s), gained points were displayed as a green area and lost points as a red area on the bar. The horizontal lines crossing the bars indicated the threshold for reaching goal A and goal B. After 15 trials, feedback for the miniblock was displayed (4s) informing the participant about the reward gained. **(B)** Summary of offer types and their effect on point count. Offers occurred with equal probability in each trial of the miniblock. Basic offers (A and B) increased either A or B points. Mixed offers (Ab and aB) added one point on one side but subtracted one point on the other side. Only accepting an offer had an effect on points. **(C)** Three different conditions modulated the difficulty to reach both thresholds by varying the number of initial points. Using an optimal agent, we chose the number of initial points, such that the agent’s probability of reaching both thresholds (G2-success) was 75% in easy, 35% in medium and 7% in hard.

Participants were presented with four different offers (A, B, Ab, and aB) that occurred with equal probability on each trial of the miniblock (see Fig 1, B). We call A or B basic offers and Ab or aB mixed offers. Accepting basic offers increased the corresponding point count, whereas accepting mixed offers transferred a single point from one scale to the other. The basic offers introduce a stochastic base rate of points, which allows participants to accumulate enough points on one or both point scales. In contrast, mixed offers allow us to identify participants’ intention to reach a state in which either both point scales are above threshold (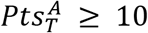 and 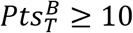) or only one point scale is above threshold (e.g. 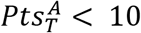 and 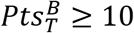; see below for more details). Rejecting an offer did not have any effect on the current point count. All participants received the same sequence of offers. We generated pseudorandomized lists for the training phase and for the three main experimental phases such that the frequency of offers reflected an equal offer occurrence probability in every list. We associated each offer with a coloured symbol to facilitate fast recognition.

Three different conditions modulated the difficulty to reach both thresholds by varying the number of initial points (Fig 1, C). We chose the number of initial points such that an optimal agent’s probability of reaching both thresholds was 75% in easy, 35% in medium and 7% in hard. The agent’s goal reaching performance for each initial point configuration was based on 10,000 simulated miniblocks with uniform offer probability (see below how we define the optimal agent). The same sequence of start conditions was presented to all participants. Pseudorandomized lists with a balanced frequency of initial point configurations were generated for the training phase and for the three main experimental phases. Note that the observed agent behaviour in the results section deviates from what we expected based on the experimental parametrization process. These discrepancies arise because we used random offer sequences (offers with equal probability) for experimental parametrization, but one specific offer sequence for the actual experiment. For example, in some miniblocks there were only few basic offers (see S1–4 Fig for details about the used offer sequence).

### Choice classification

In order to maximize reward, it was key for the participants to decide whether they should pursue the A- and B-goal in a sequential or in a parallel manner. A parallel strategy, i.e. balancing the two point scales, increases the likelihood that both goals (G2, see Table 1) will be reached at the end of the miniblock, but at the risk of failing. A sequential strategy, i.e. first secure one goal, then focus on the second one, might increase the likelihood to reach at least one goal (G1) within 15 trials, but decreases the likelihood to achieve G2.

To obtain a trial-wise measure of the pursued goal strategy, choices were classified based on the current point difference and the offer. Choices that minimized the difference between points were classified as two-goal-choice (*a_t_* = *g*2), reflecting the intention to fill both bars using a parallel strategy. Choices that maximized the difference between points were classified as one-goal-choice (*a_t_* = *g*1), reflecting the intention to pursue G1, or the intention to maintain one bar above threshold if G1-success has already been attained (see S1 Table). For example, if a participant has 8 A-points and 6 B-points and the current offer is Ab, accepting would be a g1-choice, whereas waiting would be a g2-choice. Conversely, for an aB offer, accepting would be a g2-choice and waiting a g1-choice. If the difference between points 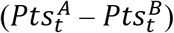 is 1 and the offer is aB, g-choice is not defined because the absolute point difference would not be changed. This also applies to the mirrored case, where the difference between points 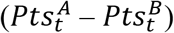 is −1 and the offer is Ab. Note that, due to the experimental design, response (accept/wait) and g-choice (g2/g1) were weakly correlated (r = 0.21). Furthermore, g-choice classification is only defined for the mixed offers (Ab and aB). The basic offers (A and B) are not informative with respect to the participants’ pursued goal strategy. Importantly, all trial-level analysis will be restricted to trials which can be related to g-choices.

### Task model

Here we will formulate the task in an explicit mathematical form, which will help us clarify what implicit assumptions we make in the behavioural model [19]. We define a miniblock of the two-goal task as a tuple

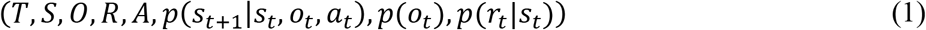

where

- *T* = 15 denotes the number of trials in a miniblock, hence *t* = 1,…, 15.
- *S* ={0,…, 20}^2^ denotes the set of task states, corresponding to the point scale of the two point types (A, and B). Hence, a state *s_t_* in trial *t* is defined as a tuple consisting of point counts along the two scales, 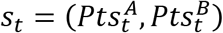.
- *o* = {*A, B, Ab, Ba*} denotes the set of four offer types, where the upper case letters denote an increase in points of a specific type and the lower case letters subtraction of points.
- *R* = {*R*_0_, *R_L_*, *R_H_*} = (0, 5,10) denotes the set of rewards.
- *A* ={0,1}denotes the set of choices, where 0 corresponds to rejecting an offer and 1 to accepting an offer.
- *p*(*s*_*t*+1_|*s_t_*, *o_t_*, *a_t_*) denotes state transitions which are implemented in a deterministic manner as *s*_*t*+1_ = *s_t_* + *a_t_* * *m*(*o_t_*), where *m*(*o_t_*) maps offer types into the point changes on the two point scales.
- 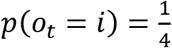 (for ∀ *i* ∈ 0) denotes a uniform distribution from which the offers are sampled.
- *P*(*r_t_*|*s_t_*) denotes the state and trial dependent reward distribution defined as

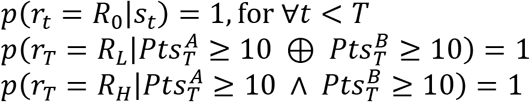 Note that in the experiment the participants are exposed to a pseudo-random sequence of offers, meaning that within one experimental block all participants observed the same sequence of offers pre-sampled from this uniform distribution (see S1–4 Fig. for additional information about the used offer sequence). For simulations and parameter estimates we use the same pseudo-random sequence of observations, hence in each trial *t* of a specific block *b* offers are selected from a predefined sequence 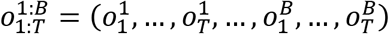, initially generated from a uniform distribution.

### Behavioural model

To build a behavioural model, we assume that participants have learned the task representation through the training session and initial instruction. Hence, the behavioural model is represented by the following tuple

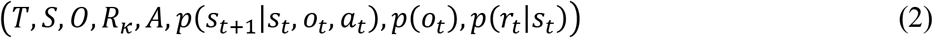

where

- *T, S, O, A, p*(*s*_*t*+1_|*s_t_, o_t_, a_t_*), *p*(*o_t_*), *p*(*r_t_*|*s_t_*) are defined the same way as in the task model.
- *R_κ_* = {0, 5, 10 · *κ*} denotes an agent-specific valuation of the rewarding states. Although the instructions for the experimental task clearly explained that participants receive a specific monetary reward depending on the final state reached during a miniblock, we considered a potential biased estimate of the ratio between G2 and G1 monetary rewards, quantified with the free model parameter *κ* ∈ [0,2]. In other words, we assumed that the participants might overestimate or underestimate the value of a G2-success, relative to a G1-success.

Importantly, the process of action selection corresponds to following a behavioural policy that maximises expected value during a single miniblock. We classified as G2-success miniblocks in which both point scales were above threshold after the final trial (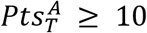 and 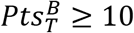). We classified as G1-success miniblocks in which only one point scale was above threshold (e.g. 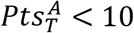 or 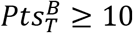).

In what follows we derive the process of estimating choice values and subsequent choices based on dynamic programming applied to a finite horizon Markov decision process ([20]; for experimental studies see also [9, 21]).

### Forward Planning

We start with a typical assumption used in reinforcement learning, namely that participants choose actions with the goal to maximize future reward. Starting from some state *s_t_* at trial *t*, offer *o_t_*, and following a behavioural policy *π* we define an expected future reward as

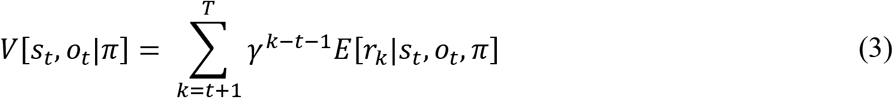

where *γ* denotes a discount rate and *E*[*r_k_* | *s_t_, o_t_, π*] denotes expected reward at some future time step *k*. The behavioural policy sets the state-action probability *π*(*a_t_*,…, *a_T_*|*s_t_*,…, *s*_*T*−1_) over the current and future trials. Hence, we can obtain the expected reward as

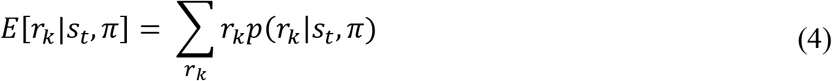

where

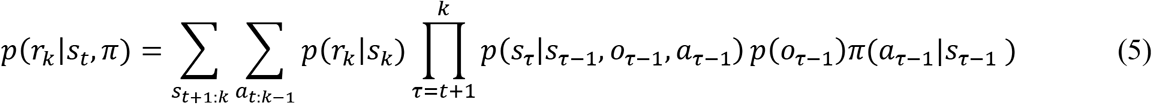

Note that we use *s*_*t*+1:*k*_, and *a*_*t*:*k*−1_ to denote a tuple of sequential variables, hence *x_m:n_* = (*x_m_*,…, *x_n_*). The key step in deriving the behavioural model was to find the policy which maximises the expected future reward, that is, the expected state-offer value. In practice, one obtains the optimal policy as

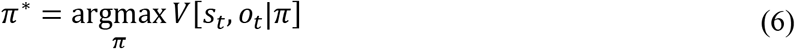

We solve the above optimization problem using the backward induction method of dynamic programming. The backward induction algorithm is defined in the following iterative steps:

i. set the value of final state *s_T_* as the reward obtained in that state 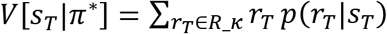
ii. compute state-offer-action value as 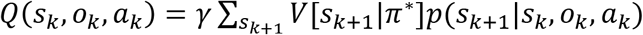
iii. set optimal choice for given state-offer pair as 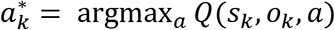
iv. define the expected value of state *s_k_* under optimal policy *π*\* as 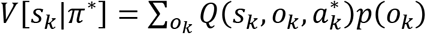
v. repeat steps (ii) – (iv) until *k* = *t*

Hence, for a fixed value of the reward ratio (*κ*) an optimal choice at trial *t* corresponds to

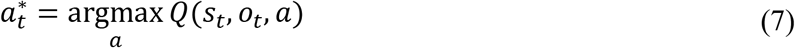

We will define the optimal agent as an agent who has a correct representation of the reward ratio (*κ* = 1) and does not discount future reward (*γ* = 1). We illustrate in Fig 2 the Q-value to accept, estimated for the case of the optimal agent in an example trial 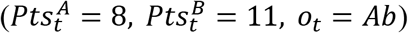.

**Fig 2.**
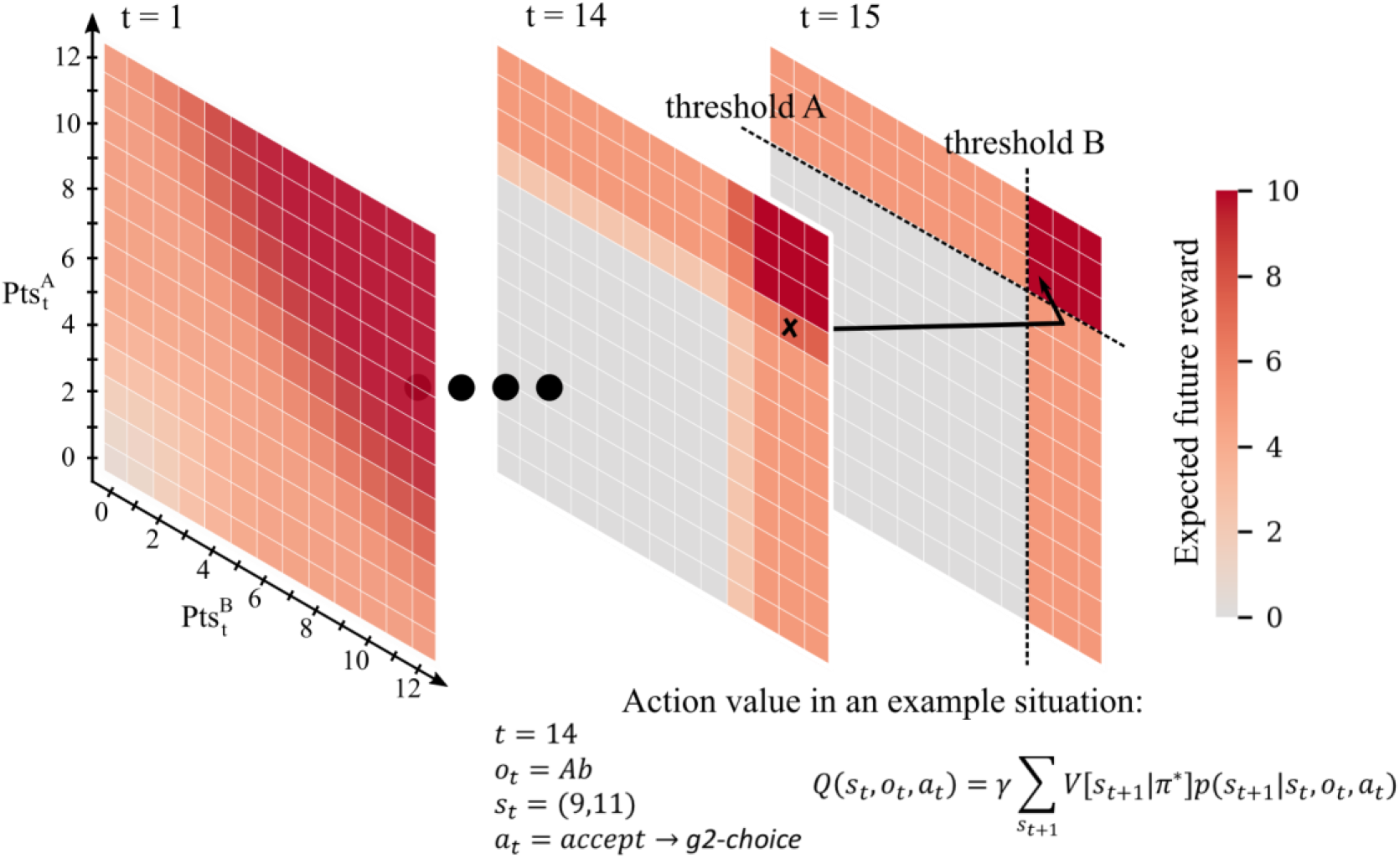
Illustration of the state space and associated expected future reward for the optimal agent (*γ* = 1, *κ* = 1). The black arrow shows a hypothetical transition in the state space. In trial 14 the participant has 9 A-points and 11 B-points (marked by the black cross) and accepts an offer Ab, gaining one A-point and losing one B-point (g2-choice). In the resulting state, both thresholds are reached; thus, the value of that state is 10 Cents. Similarly, the action that leads to that state has an associated Q-value of 10 Cents. In this example the agent would just have to wait in the last trial (15) to gain a 10 cents reward.

### Response likelihood

Participants might compute expected values by mentally simulating and comparing sequences of actions towards the end of the miniblock. To illustrate the benefits of planning we consider the following example: There are 3 trials left in the current miniblock, and the participant has 9 A-points and 9 B-points (10 is threshold), and she receives offer Ab. Planning would, for example, allow to compute the probabilities for G2 when choosing either wait or accept. By waiting the participant would enter the second last trial with 9 A-points and 9 B-points. Receiving offer A or B in the second last trial (0.5 probability) followed by the complementary offer A or B in the last trial (0.25 probability) would grant G2. When choosing accept, the participant will have in the second last trial 10 A-points and 8 B-points. Consequently, she would need two consecutive B-offers (0.25 *0.25 probability) to achieve G2. Hence, by planning ahead one would conclude that wait gives the highest probability for a G2-success.

Still, planning an arbitrary number of future steps is complex and unrealistic. Hence, we make an assumption that the process of optimal action selection described above is perturbed by noise (planning noise, and response noise) which we quantify in the form of a parameter *β*, denoting response precision. Hence, this precision parameter is critical to characterize the participants’ reliance on forward planning. Since the difference in expected future rewards of a g1- or g2-choice is high when the goal is close (S5 Fig), *β* is able to selectively capture g-choice performance at the end of the miniblock. Furthermore, instead of an elaborate planning process participants might use a simpler heuristic when deciding which action to select. We capture this heuristic in form of an additional offer-state-action function *h*(*o_t_, s_t_, a_t_, θ*) which evaluates choices relative to possible goals. We describe this heuristic evaluation below. Overall, we can express the response likelihood (the probability that a participant makes choice *a_t_*) as

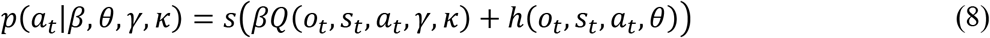

where *s*(*x*) denotes the softmax function.

### Choice heuristic

The choice heuristic is defined relative to the current offer *o_t_*, current state *s_t_*, and possible choices *a_t_*. Importantly, we will interpret the choice heuristic in terms of participants’ biases towards approaching both goals in a sequential or parallel manner. Hence, it is more intuitive to define the choice heuristic as choice biases relative to the goals, and not accept-reject choices. The choice heuristic is defined as follows

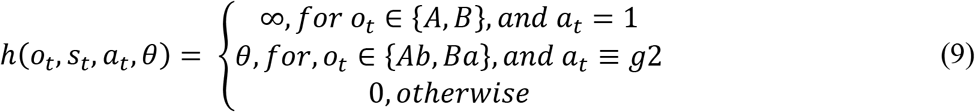

where *a_t_* ≡ *g*2 denotes choices (accept or reject) which can be classified as g2-choices (see subsection Choice classification for details). In summary, a choice which reduces the point difference 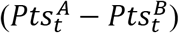, for the given offer and the current state, is classified as g2-choice and choice which increases the point difference as g1-choice. Essentially, the strategy preference parameter *θ* reflects participants’ preference for pursuing a sequential (negative values) or parallel (positive values) strategy. For example, some participants might have a general tendency to pursue goals in a parallel manner, independent of the actual *Q*-values. Conversely, participants may prefer a more cautious sequential approach. Note that we expected this parameter to make the most significant contribution to participants’ deviation from optimal behaviour, reflecting their reliance on decision heuristics early in the miniblock.

Finally, for those choices which can be classified as g2- or g1-choices, we can express the response likelihood in a simplified form, in terms of free model parameters *β, θ, γ, κ* (Table 2). We refer to the difference between Q-values for g-choice as the differential expected value (*DEV*),

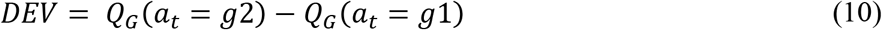

**Table 2.**
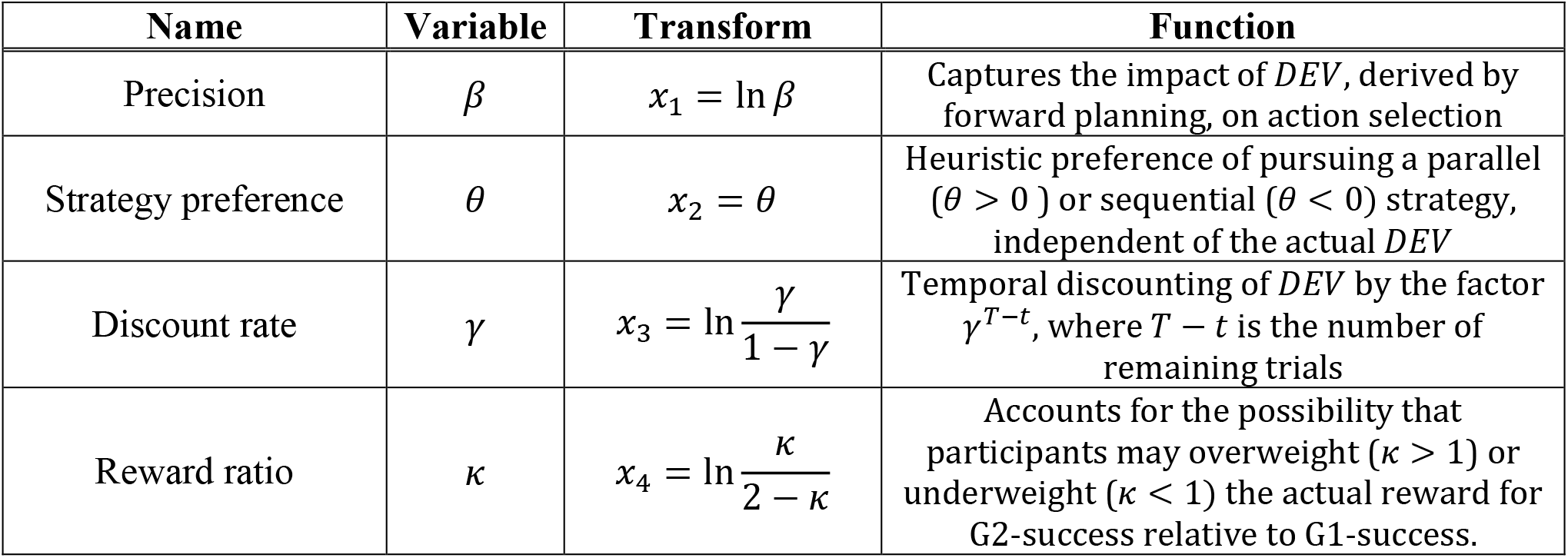
Summary of four free model parameters, the variables, the transformations used to map values to unconstrained space and their function in modelling participant behaviour.

Using *DEV*, we defined the probability of making a g2-choice as

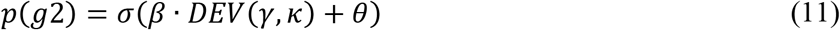

where 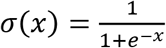 denotes the logistic function. Note that the probability of g1-choice becomes *p*(*g*1) = 1 − *p*(*g*2).

### Optimal agent comparison and general data analysis

We compared participant behaviour with simulated behaviour of an optimal agent. To summarize, we denote the optimal agent as the agent which has a correct representation of the reward function (*κ* = 1), does not discount future rewards (*γ* = 1), is not biased in favour of any choice (*θ* = 0), and who generates deterministic g-choices based on *DEV*-values (corresponding to *β* → ∞ in the response likelihood, that is, the argmax operator). The optimal agent deterministically accepts A and B offers.

When simulating agent behaviour to evaluate successful goal reaching, the agent received the same sequence of offers and initial conditions as the participants. Analysis on the level of g-choices was performed by registering instances in which the g-choice of a participant differed from the g-choice the optimal agent would have made in the same context 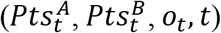. Trials with A or B offers and trials in which G2 had already been reached, were excluded from the g-choice analysis.

The goal of this comparison between summary measures of both optimal agent and participants was two-fold: First, we used this comparison to visualize deviations from optimality and motivate the model-based analysis which was used to test the hypothesis that a shift from heuristics to forward planning may explain these deviations. Second, plotting suboptimal g-choices instead of g-choices (Fig. 4) makes behaviour between participants more comparable. Plotting the proportion of g-choices averaged across participants would have been mostly uninformative because the significance of a g-choice depends on the current state, which is a consequence of the individual history of past choices within a miniblock. By registering deviations from an optimal reference point, we circumvent this state dependence of g-choices.

We used a sign test as implemented in the “sign_test” function of python’s “Statsmodels”[22] package to test whether participants total reward and success rates differed significantly from the optimal agent’s deterministic performance. We reported the p-value and the m-value *m* = (*N*(+) − *N*(−))/2, where *N*(+) is the number of values above 0 and *N*(−) is the number of values below and. To test for learning effects (in the main experimental phase), we used mixed effects models as implemented in R [23] with the “lm4” package [24]. Intercepts and slopes were allowed to vary between participants. p-values were obtained using the “lmerTest” package [25].

### Hierarchical Bayesian data analysis

To estimate the free model parameters (Table 2) that best match the behaviour of each participant, we applied an approximate probabilistic inference scheme over a hierarchical parametric model, so-called stochastic variational inference (SVI) [26].

As a first step, we define a generic (weakly informative) hierarchical prior over unconstrained space of model parameters. In Table 2 we summarize the roles of free model parameters of our behavioural model and the corresponding transforms that we used to map parameters into an unconstrained space. We use ***x***^*n*^ to denote a vector of free and unconstrained model parameters corresponding to the *n*th participant. Similarly, ***μ*** and ***σ*** will denote hyperpriors over group mean and variance for each free model parameter. We can express the hierarchical prior in the following form

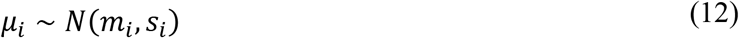

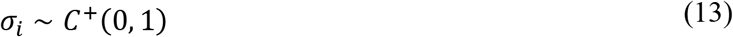

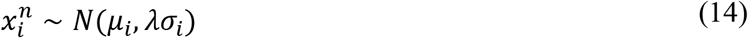

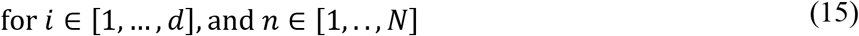

where *C*^+^(0,1) denotes a Half-Cauchy prior with scale *s* = 1, *d* number of parameters, and *N* number of participants. Note that by using this form of a hierarchical prior we make an explicit assumption that parameters defining the behaviour of each participant are centred on the same mean and share the same prior uncertainty. Hence, both the prior mean and uncertainty for each parameter are defined at the group level. Furthermore, the hyper-parameters of the prior *η* = (*m*_1_,…, *m*_4_, *s*_1_,…, *s*_4_, *λ*) are also estimated from the data (Empirical Bayes procedure) in parallel to the posterior estimates of latent variables *θ* = (*μ*_1_,…, *μ*_4_, *σ*_1_,…, *σ*_4_, ***x***^1^,…, ***x***^*N*^). For more details, see supporting information (S1 Notebook).

The behavioural model introduced above defines the response likelihood, that is, the probability of observing measured responses when sampling responses from the model, condition on the set of model parameters (***x***^1^,…,***x***^*N*^). The response likelihood can be simply expressed as a product of response probabilities over all measured responses *A* = (***a***^1^,…, ***a***^*N*^), presented offers *O* = (***o***^1^,…, ***o***^*N*^), and states (point configurations) visited by each participant *S* = (***s***^1^,…, ***s***^*N*^) over the whole experiment

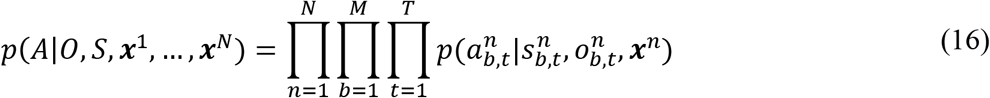

where *b* denotes experimental block, and *t* a specific trial within the block.

To estimate the posterior distribution (per participant) over free model parameters, we applied the following approximation to the true posterior

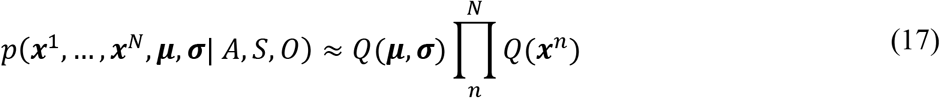

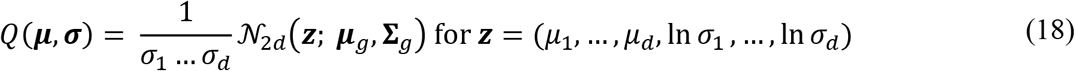

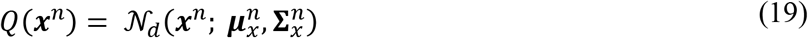

Note that the approximate posterior captures posterior dependencies between free model parameters (in the true posterior) on both levels of the hierarchy using the multivariate normal and multivariate log-normal distributions. However, for practical reasons, we assume statistical independence between different levels of the hierarchy, and between participants. Independence between participants is justified by the structure of both response likelihood (responses are modelled as independent and identically distributed samples from conditional likelihood) and hierarchical prior (a priori statistical independence between model parameters for each participant).

Finally, to find the best approximation of the true posterior given the functional constraints of our approximate posterior, we minimized the variational free energy F[Q] with respect to the parameters of the approximate posterior.

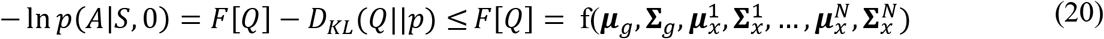

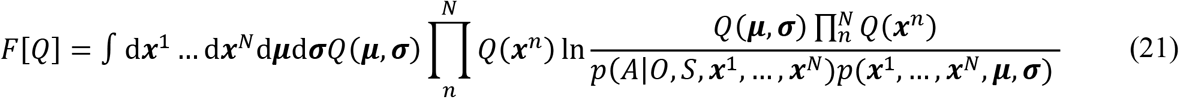

The optimization of the variational free energy F[Q] is based on the SVI implemented in the probabilistic programing language Pyro [27] and the automatic differentiation module of PyTorch [28], an open source deep learning platform.

As a final remark, we would like to point out that it is possible to use a different hierarchical prior [29], different parametrization of the hierarchical model [30] or different factorization of the approximate posterior (e.g., mean-field approximation). However, through extensive comparison of posterior estimates on simulated data, we have determined that the presented hierarchical model and the corresponding approximate posterior provide the best posterior estimate of free model parameters among the set of parametric models we tested (S1 Notebook).

## Results

To investigate how the balance between computationally costly forward planning and heuristic preferences changes as a function of temporal distance from the goals, participants performed sequences of actions in a novel sequential decision-making task. The task employed a two-goal setting, where participants had to decide between approaching the two goals in a sequential or in a parallel manner. We first performed a standard behavioural analysis, followed by a model-based approach showing that participants use a mixture of strategy preference and forward planning to select their action.

### Standard behavioural analysis

We first analysed the general performance of all participants and – for each miniblock and trial – compared it to the behaviour of an optimal agent possessing perfect knowledge of the task and performing full forward planning to derive an optimal policy that maximizes total reward. The motivation of this comparison was to detect differences between how the optimal agent and participants perform the task. These differences will motivate our model-based analysis below. To compute and compare optimal vs individual policies, all participants and the agent received exactly the same sequence of offers and start conditions. The difference in total reward between participants and agent was significant (m = −35.5, *p* < 0.001), where participants earned 388.5 Cents (SD = 13.6) and the agent earned 405 Cents. As expected, both participants and agent earned more money in the easy condition than in the medium condition and least in the hard condition (Fig 3, A, C). In the easy and medium condition, the agent earned significantly more than the participants (easy: M = 8.7 Cents, SD = 8.4, m = −33, *p* < 0.001; medium: M = 7.2 Cents, SD = 7.0, m = −30, *p* < 0.001). In the hard condition, the total reward did not differ significantly between the participants and agent, m = 0.5, p > 0.99 (Fig 3, E). These results show that participant performance was generally close to the optimal agent but differed significantly in the easy and medium condition.

**Fig 3.**
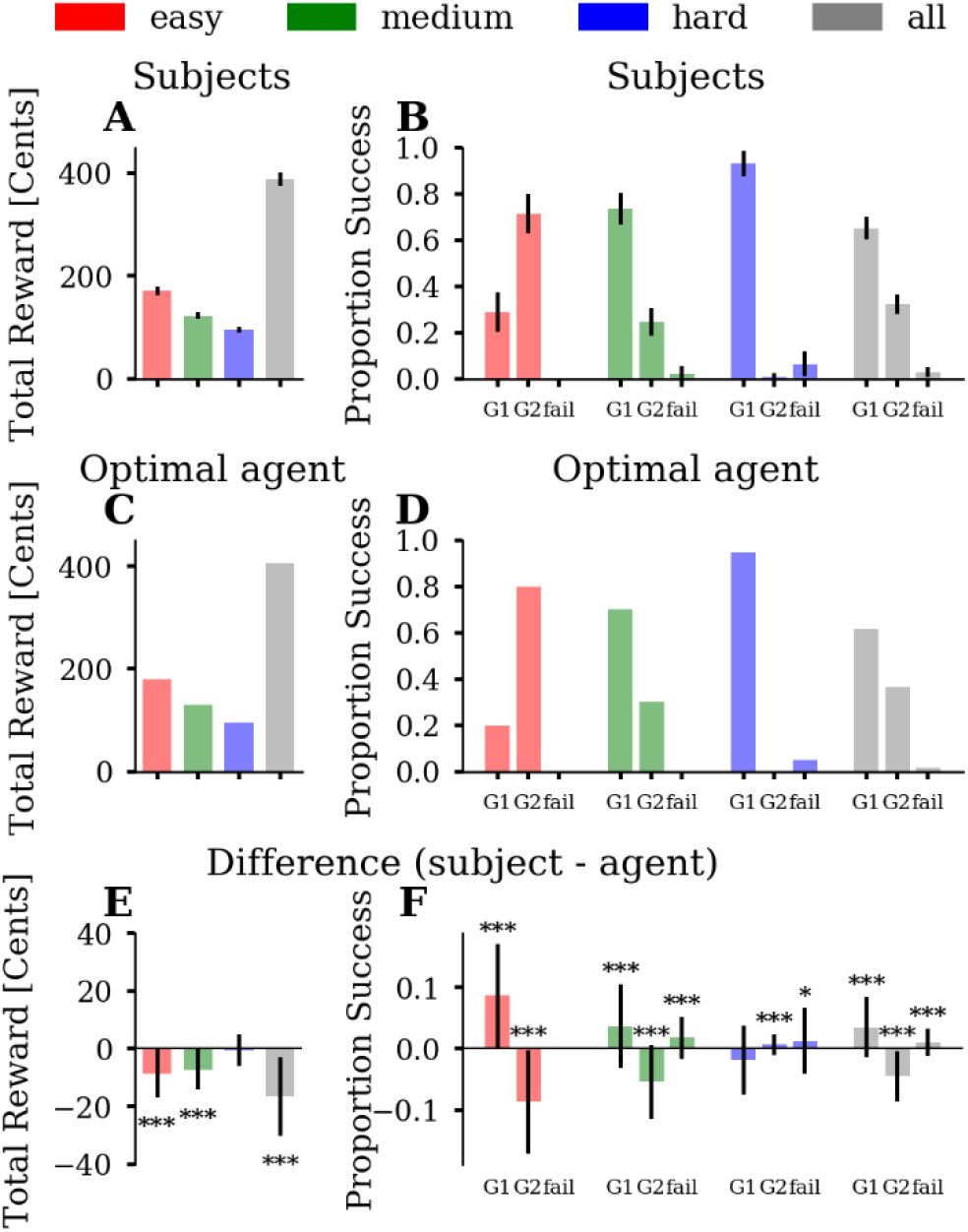
Standard analyses of total reward and comparison to the optimal agent. **(A)** Average total reward across participants. The three conditions are colour-coded (easy = red, medium = green, blue = hard) and the average over conditions is shown in grey. Error bars depict the standard deviation (SD). **(B)** Proportion of successful goal-reaching averaged across participants, for each of the three conditions. We plot the proportion of reaching, at the end of a miniblock, a single goal (G1), both goals (G2), or no goal (fail). The fourth block of bars in grey represents the proportions averaged over all three conditions. Error bars depict SD. **(C)** Simulated total reward of the optimal agent. **(D)** The goal-reaching proportions of the optimal agent. **(E)** Average difference between participants and agent with error bars depicting SD. **(F)** Averaged difference of proportion success between participants and agent with error bars depicting SD. One can see that the average goal-reaching proportions of participants were close to the agent’s proportions. However, participants, on average, reached G2 less often than the agent. Asterisks indicate differences significantly greater than zero (Sign-test, * ≙ p < 0.05, ** ≙ p < 0.01, *** ≙ p < 0.001).

Next, we analysed participants’ goal reaching success and compared it to the optimal agent. There were three possible outcomes in a miniblock: Achieving G1 (goal A or B), achieving G2 (A & B) or fail (neither A nor B). The main experiment comprised 20 miniblocks of each difficulty level modulating difficulty to reach G2. As expected, participants reached on average G2 more often in the easy (M = 71%, SD = 8%) than in the medium condition (M = 25%, SD = 6%), m = 44.5, p < 0.001. In the hard condition, participants reached G2 in only 1% (SD = 2%) of the miniblocks. Participants failed to reach any goal in 2% (SD = 3%) of the miniblocks in the medium and in 6 % (SD = 5%) of the miniblocks in the hard condition. They never failed in the easy condition (Fig 3, B). The agent reached G2 in 80% in the easy, in 30% in the medium and in 0% in the hard condition (Fig 3, D). Note that G2 cannot be reached in all miniblocks. We simulated all possible choice sequences (n = 2^15) for a given miniblock and evaluated whether G2 was theoretically possible. According to these simulations, 90% G2 performance can be reached in the easy, 35% in the medium and 5% in the hard condition.

When comparing participants’ goal reaching success with the agent, we found that, on average, there was a consistent pattern of deviations in the easy and medium conditions (Fig 3, F). In the easy condition, participants reached G2 on average 9% (SD = 8%) less often than the agent (m = −33, p < 0.001), but reached G1 9% (SD = 8%) more often (m = 33, p < 0.001). In the medium condition, participants reached G2 on average 6% (SD = 6%) less often than the agent (m = −26, p < 0.001) but reached G1 4% (SD = 7%) more often (m = 16.5, p < 0.001). While the agent never failed, participants had a 2% (SD = 3%) fail rate (m = 11.5, p < 0.001). In the hard condition, participants reached G2 on average 0.6% (SD = 1.6%) more often than the agent (m = 5.5, p < 0.001). G1 (m = - 7, p = 0.087) and fail-rate (m = 3.5, p = 0.42) did not differs significantly between participants and agent. In summary, these differences in successful goal reaching between participants and the agent explains the difference in accumulated total reward: Participants obtained less reward than the agent because on average they missed some of the opportunities to reach G2 in the easy and medium condition and sometimes even failed to achieve any goal in the medium and hard condition.

How can these differences in goal-reaching success be explained? To address this, we used the mixed-offer trials to identify which strategy a participant was pursuing in a given trial and compared the strategy choice to what the agent would have done in this trial. We classified strategy choices as evidence either of a parallel or a sequential strategy. With the parallel strategy (g2), participants make choices to pursue both goals in a parallel manner, while with a sequential strategy (g1), participants make choices to reach first a single goal and then the other. We inferred that participants used a g2-choice for a specific mixed-offer trial when the difference between the points of the two bars was minimized, while we inferred a g1-choice when the difference between points was maximized (see Methods). We categorized a participant’s g2-choice as suboptimal when the optimal agent would have made a g1-choice in a specific trial and vice versa. Fig 4, A-D shows the proportions of suboptimal g-choices in mixed-offer trials. In the easy condition, participants made barely any suboptimal g2-choice (mean = 0%, SD = 0.001%), but 29% (SD = 10%) suboptimal g1-choices (Fig 4, A). This means that participants, on average, preferred a sequential strategy more often than would have been optimal. In the medium condition participants made on average 6% (SD = 3%) suboptimal g2-choices and 28% (SD = 11%) suboptimal g1-choices. Similar to the easy condition, participants, on average, preferred a sequential strategy where a parallel strategy would have been optimal. In the hard condition, this pattern reversed. Participants made on average 40% (SD = 12%) suboptimal g2-choices, relative to the agent, and 11% (SD = 6%) suboptimal g1-choices. Participants’ suboptimal g-choices were also reflected in goal reaching success. In the easy and medium condition, suboptimal g1-choices, relative to the agent, resulted in a higher proportion of reaching G1, and a lower proportion of reaching G2. In the hard condition, suboptimal g2-choices led to occasional fails and a tiny margin of reaching G2. However, despite suboptimal g2-choices, participants still reached G1 in 93% (SD = 6%) of the miniblocks.

**Fig 4.**
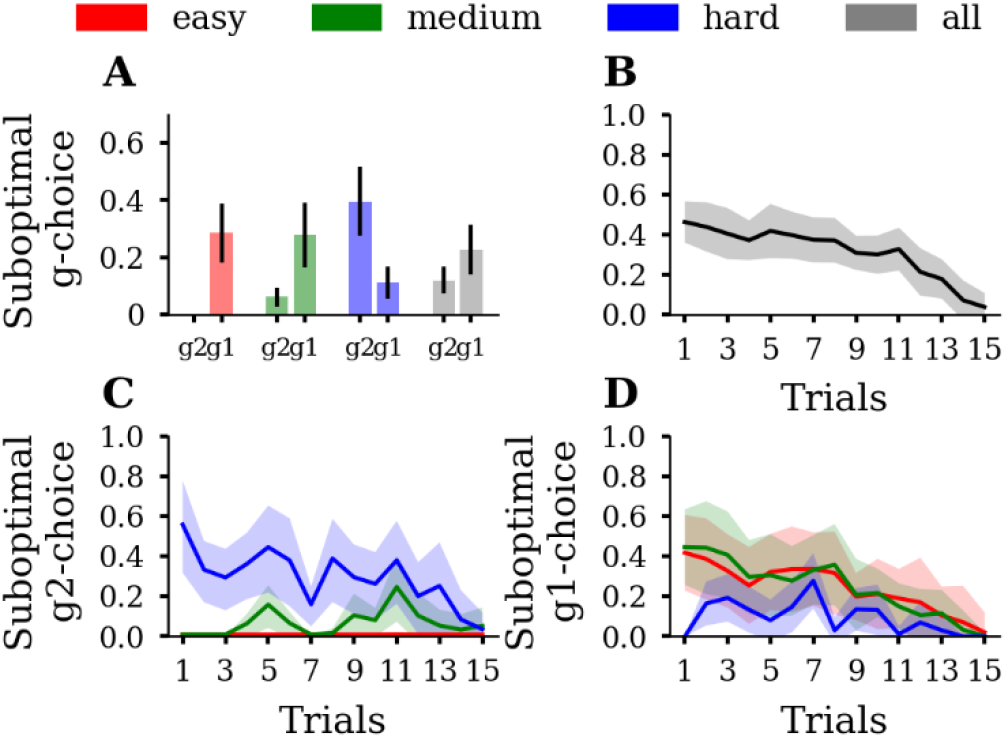
**(A)** Proportions of suboptimal g1-choices (g1) and suboptimal g2-choices (g2), averaged over participants. Participants tend to make suboptimal g1-choices in the easy and medium condition while this pattern reverses in the hard condition. Error bars depict SD. Conditions are colour coded. **(B)** Suboptimal g-choices as a function of trial averaged over participants. Shaded areas depict SD. **(C)** Suboptimal g2-choices as a function of trial averaged over participants. **(D)** Suboptimal g1-choices as a function of trial averaged over participants. In both C and D, one can see that participants made more suboptimal g-choices at the beginning of the miniblock than close to the final trial. Shaded areas depict SD.

As the first test of our prediction that participants tend to use more forward planning when temporally proximal to the goal, we analysed suboptimal decisions as a function of trial time. As expected, suboptimal decisions, relative to the agent, decreased over trial time (Fig 4, B). While in the first trial, 42% (SD = 19%) of participants’ g-choices deviated from the agent’s g-choices, participant behaviour converged to almost optimal performance towards the end of the miniblock, with only 4% deviating g-choices (SD = 7%). We also simulated a random agent that accepts all basic A or B offers but guesses on mixed offers (S6–7 Fig). S7 Fig B shows that the random agent makes approximately 50 % suboptimal g-choices across all trials in the miniblock. That means participants used non-random response strategies, i.e. planning or heuristics, since their pattern of suboptimality across trials deviated from the straight-line pattern of the random agent.

In the hard condition, the number of suboptimal g2-choices similarly decreased, but not in the easy and medium condition (Fig 4, C). The number of suboptimal g1-choices decreased across trials in the easy and medium, but not in hard condition (Fig 4, D). Note that in easy and the medium conditions, opportunities to make suboptimal g2-choices are generally scarce, because the difference between action values *DEV* = *Q_G_*(*g*2) – *Q_G_*(*g*1) was mostly positive, which means that a g2-choice was mostly optimal. Similarly, in the hard condition, as there was a low number of opportunities to make suboptimal g1-choices, there was no clear decrease in the number of suboptimal g1-choices.

Although these findings of diminishing suboptimal choices over the course of miniblocks may be explained by the participants’ initial employment of a suboptimal heuristic, there is an alternative explanation because we used an optimal agent, which uses a max operator to select its action: If this agent computes, by using forward planning, a tiny advantage in expected reward of one action over the other, the agent will always choose in a deterministic fashion the action with the slightly higher expected reward. Therefore, at the beginning of the miniblock, where the distance to the final trial is largest, the difference between goal choice values *DEV* = *Q_G_*(*g*2) – *Q_G_*(*g*1) (S5 Fig) is close to 0. The reason for this is that a single g2-choice at the beginning of the miniblock does not increase the probability for G2-success by much. However, when only few trials are left, a single g2-choice might make the difference between winning or losing G2. Since *DEVs* are close to 0 at the initial trials we cannot exclude the possibility yet that participants actually may have used optimal forward planning just like the agent but did not use a max operator. Instead, participants may have sampled an action according to the computed probabilities of each action to reach the greater reward in the final trial. Such a sampling procedure to select actions would also explain the observed pattern of diminishing suboptimal g-choices over the miniblock (Fig. 4 B-C). To answer the question, whether there is actually evidence that participants use heuristics, when far from the goal, even in the presence of probabilistic action selection of participants, we now turn to a model-based analysis.

### Model-based behavioural analysis

To infer the contributions of participants’ forward planning and heuristic preferences, we conducted a model-based analysis. If we find that participants’ strategy preference *θ* is smaller or larger than zero, we can conclude that participants indeed used a heuristic component to complement any forward planning. This is especially relevant for choices early in the miniblock as *DEV* values are typically close to zero. Indeed, when inferring the four parameters for all 89 participants using hierarchical Bayesian inference, we found that participants’ g-choices were influenced by a heuristic strategy preference in addition to a forward planning component (Fig 5, A). For 74 out of 89 participants, we found that the 90% credibility interval (CI) of the posterior over strategy preference did not include zero. 68 of these participants had a positive strategy preference, meaning they preferred an overall strategy of pursuing both goals in parallel. Six of these participants had a negative strategy preference, meaning they preferred to pursue both goals sequentially. The median group hyperparameter of strategy preference was 0.55 (90% CI = [0.47, 0.63]). For example, a participant with this median strategy preference, in a mixed-offer trial where *DEV* = 0, would make a g2-choice with 63% probability, whereas a participant without a strategy preference bias, i.e. *θ* = 0, would make a g2-choice with 50% probability. After the experiment, we had asked participants whether they used any specific strategies to solve the task and to give a verbal description of the used strategy. Reports reflected three main patterns: Pursuing one goal after the other (sequential strategy), promoting both goals in a balanced way (parallel strategy), and switching between sequential and parallel strategy, depending on context (mixed strategy). Reported strategies are in good qualitative agreement with the estimated strategy preference parameter (S8 Fig), supporting our interpretation of this parameter. Notably, the task instructions, given to the participants prior to the experiment, did not point to any specific heuristic (S1 Text). Altogether, the non-zero strategy preference in 83% of participants indicates that suboptimal decisions within a miniblock (see Fig 4) are not only caused by probabilistic sampling for action selection, but also by the use of a heuristic strategy preference.

**Fig 5.**
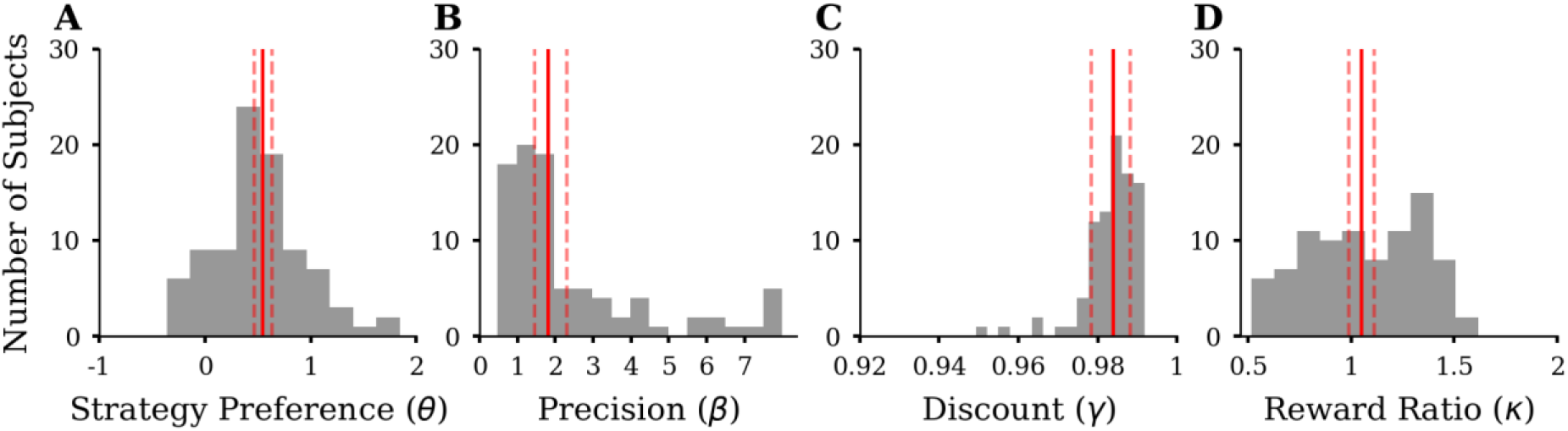
Summary of inferred parameters of the four-parameter model for all 89 participants. We show histograms of the median of the posterior distribution, for each participant. Solid red lines indicate the median of the group hyperparameter posterior estimate with dashed lines indicating 90% credibility intervals (CI). **(A)** Histogram of strategy preference parameter *θ*. **(B)** Histogram of precision parameter *β* (last bin containing values > 8). **(C)** Histogram of discount parameter *γ*. **(D)** Histogram of reward ratio parameter *κ*.

As expected, we found that the *DEV* (see Table 1) derived by forward planning influenced action selection (median group hyperparameter of the inferred precision *β* = 1.82, 90% CI = [1.45, 2.3], Fig 5, B). For example, a hypothetical participant with parameters similar to the group hyperparameters (0 = 0.55 and *β* = 1.82), when encountering a *DEV* = 0.5, would make a g2-choice with 82% probability. Increasing *DEV* by 1 would increase the g2-choice probability to 96%. In contrast, a participant with low precision but the same median strategy preference (0 = 0.55 and *β* = 0.5), when encountering a *DEV* = 0.5, would make a g2-choice with 69% probability. Increasing *DEV* by 1 would increase g2-choice probability to 79%. We found evidence only for weak discounting of future rewards, as for most participants the inferred discount was close to 1 (median of the inferred discount parameter *γ* = 0.984, 90% CI = [0.978, 0.988], Fig 5, C). We found that some participants used a reward ratio different from the objective value of 1 (CI not containing 1). Twelve participants had a reward ratio greater than 1 and 17 participants had a reward ratio smaller than 1. However, the median group hyperparameter of the inferred reward ratio was close to the objective value of 1 (*κ* = 1.05, 90% CI = [0.99, 1.11], Fig 5, D). A reward ratio of 1.2 means, that participants behaved as if the value of achieving G2 would be 2.4 times the value of achieving G1(when in reality the reward is only double as high). While strategy preference has its greatest influence during the first few trials of a miniblock, the reward ratio has an influence only when forward planning, i.e. changes the *DEV*, and will therefore affect action selection most during the final trials of a miniblock. In addition, we found only low posterior correlation between the strategy preference and reward ratio parameter, indicating that these two parameters model distinct influences on goal reaching behaviour.

To show that our model with constant parameters is able to capture a dynamic shift from heuristic decision making to forward planning we conducted two sets of simulations where we systematically varied the response precision *β* and the strategy preference parameter *θ*. First, we simulated behaviour where we varied β between 0.25 and 3 with θ, *γ*, and *κ* sampled from their fitted population mean (S1-2 Movie). S2 Movie, B shows that the higher *β*, the fewer suboptimal g-choices are made towards the end of the miniblock. Second, we simulated behaviour where we varied *θ* varied between −1 and 1 with *β*, *γ*, and *κ* sampled from their fitted population mean (S3-4 Movie). S4 Movie, B shows that a change in *θ* affects the number of suboptimal g-choices made at the beginning but not at the end of the miniblock. These two results support the argument that the *θ* parameter is able to capture heuristic decision making at the beginning of the miniblock while the *β* parameter is able to capture planning behaviour at the end of the miniblock. The reason for this interaction between parameter effect and trial number is that differential expected value (*DEV*) computed by forward planning is close to zero at the beginning of the miniblock but increases towards the end of the miniblock (S5 Fig). For small *DEVs*, the influence of *β* on choice probability is marginal; therefore, the relative influence of the strategy preference parameter *θ* is high, and behaviour is explained by using the heuristic. For higher trial numbers, i.e. closer to the end of the miniblock, *DEVs* tend to be high so that the influence of the response precision *β* is high, and the relative influence of *θ* is low; therefore, towards the end of the miniblock behaviour is explained by forward planning with a shift in between, depending on the dynamics of the *DEV*. We also implemented a model with changing parameters over trials and compared it to the constant model. Parameters were fit separately for three partitions of the miniblock, i.e. early (trials 1-5), middle (trials 6-10) and late trials (11-15). Model comparisons showed that this model with changing parameters had lower model evidence compared to the model with constant parameters (S9 Fig). We interpret these results as further evidence that the described constant parameterization is sufficient to describe a hidden shift from using a heuristics to forward planning.

Finally, as an additional test of the hypothesis that participants rely more on heuristic preferences when the goal is temporally distant, we conducted a multiple regression analysis (Fig 6, A). To do this, we divided the data into the first (first 7 trials) and the second half (last 8 trials) of miniblocks, and computed, for each participant the proportion of g2-choices in the mixed-offer trials. We fitted, across participants, these proportions of g2-choices against 6 regressors: strategy preference, precision, discount rate, reward ratio, a dummy variable coding for the first and second miniblock half and interaction between strategy preference and miniblock half. We found a significant interaction between strategy preference and miniblock-half (p < 0.001), demonstrating that strategy preference is more predictive for the proportion of g2-choices in the first half of the miniblock than in the second half. Fig 6, B visualizes the interaction effect showing that the slope of the marginal regression line for the first half of the miniblock is greater than the slope of the marginal regression line for the second half of the miniblock. This finding provides additional evidence that participants rely on heuristic preferences when the goal is temporally far away but use differential expected values (*DEV*) derived by forward planning when the goal is closer.

**Fig 6.**
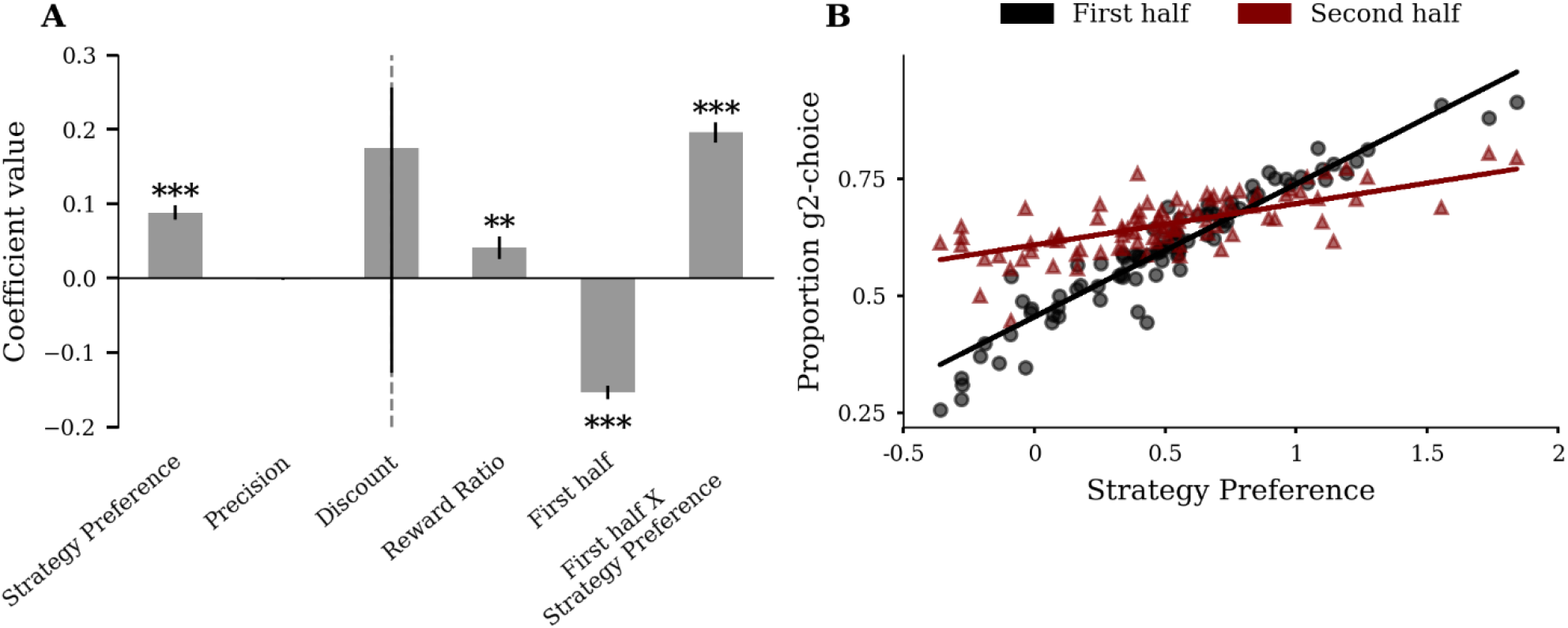
Strategy preference is more predictive for participant’s proportion of g2-choices in the first than in the second half of the miniblock. **(A)** Linear regression of proportion g2-choice against parameters from the four-parameter model, a dummy variable coding for miniblock-half and interaction between miniblock-half and strategy preference. The significant interaction term supports the hypothesis that the influence of strategy preference on g2-choice proportion is greater in the first than in the second half of the miniblock. Error bars represent SE. Asterisks indicate coefficients significantly different from 0 (t-test, * ≙ p < 0.05, ** ≙ p < 0.01, *** ≙ p < 0.001). **(B)** Strategy preference plotted against the proportion of g2-choices in the first half of the miniblock (black) and in the second half of the miniblock (red). Solid lines represent marginal regression lines.

In addition, we conducted model comparisons, posterior predictive checks and parameter recovery simulations to test whether our model is an accurate and parsimonious fit to the data. First, we compared variants of our model, where we fixed individual parameters (S9 Fig). Adding *θ* and *β*increased model evidence, confirming their importance in explaining participant behaviour. The three-parameter model (*θ, β, κ*) had the highest model evidence among all 16 models. Adding *γ* did not increase model evidence. This result is consistent since we found only little evidence for discounting when fitting the parameters, see Fig. 5C. To test whether participants used condition-specific response strategies (e.g., use heuristics in the easy and hard but plan forward in the medium difficult condition) we estimated model parameters separately for conditions. However, the condition-wise model had lower model evidence compared to the conjoint model, indicating that participants use a condition-general approach to arbitrate between using a heuristic and planning ahead. Second, we simulated data using the group mean parameters as inferred from the participants’ data and compared it to the observed data. Visual inspection shows that both the simulated performance pattern (S10 Fig) and the simulated frequency of suboptimal g-choices (S11 Fig) closely resemble the experimentally observed patterns (Fig. 3 and 4). Third, we simulated data using participants’ posterior mean and tested whether we could reliably infer parameters (S1 Notebook). Results showed that the inferred *β, θ* and *κ* align with the true parameter value, but simulation-based calibration [31] suggests that estimates of *γ* are biased. Taken together, our model provides a good fit to the data, where the data are informative about the three parameters *β, θ* and *κ*.

We also tested whether participants showed learning effects in the main experimental phase. In a first linear model, the depended variable was the total reward and the predictor was the experimental block number (miniblock 1-20, miniblock 21-40, miniblock 41-60). The analysis revealed a significant but small main effect of experiment block (*β* = 5.4, SE = 0.5, p < 0.001). In a second logistic model the dependent variable was suboptimal goal choice (1 = suboptimal, 0 = optimal) and the predictor was experiment block. The second analysis revealed a significant but small main effect of experiment block on the probability to make a suboptimal g-choice (*β* = −0.084, SE = 0.02, p < 0.001). Furthermore, we fitted the three parameter model (*θ, β, κ*) separately for experiment blocks. Model comparisons revealed that the experiment block-wise model had lower model evidence compared to the conjoint model (S9 Fig.).

As a final control analysis, we used logistic regression to establish how the absolute difference between A- and B-points affects goal choice as a function of the number of trials remaining in the miniblock. If participants rely on a fixed strategy preference when far from the goal, there should be no effect of absolute score difference on goal choice at the start of miniblocks. In this model the depended variable was goal choice (1 = g2, 0 = g1) and the predictors were absolute score difference 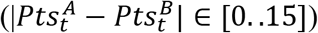, miniblock-half (1 = trial 1-7, 0 = trial 8-15) and the interaction term absolute score difference*miniblock-half. There was a significant main effect of absolute score difference (*β* = 0.14, SE = 0.008, p < 0.001) and miniblock-half (*β* = 0.29, SE = 0.039, p < 0.001). Importantly, the analysis revealed a significant interaction between miniblock-half and absolute score difference (*β* = −0.2, SE = 0.013, p < 0.001). This means that goal choice was more affected by the absolute score difference in the second half the miniblock compared to the first half. The analysis supports our conclusion that participants relied on a heuristic strategy preference when far from the goal.

## Discussion

In the current study, we investigated how humans change the way they decide what goal to pursue while approaching two potential goals. To emulate real life temporally extended decision making scenarios of goal pursuit, we used a novel sequential decision making task. In this task environment, decisions of participants had deterministic consequences, but the options given to participants on each of the 15 trials were stochastic. This meant that especially during the first few trials, participants could not predict with certainty what goal was achievable. Using model-based analysis of behavioural data we find that most participants, during the initial trials, relied on computationally inexpensive heuristics and switched to forward planning only when closer to the final trial.

We inferred the transition from a heuristic action selection to action selection based on forward planning using a model parameter that captured participants’ preference for pursuing both goals either in a sequential or parallel manner. This strategy preference had its strongest impact for the first few trials, when participants, due to the stochasticity of future offers, could not predict well which of the two available actions in a mixed trial would enable them to maximize their gain. This can be seen from Eq. 11 where two terms contribute to making a decision: the term containing the differential expected value (*DEV*) and the strategy preference *θ*. In our computational model, the *DEV* is the difference between the expected value of a sequential strategy choice and a parallel strategy choice. The *DEV* enables the agent to choose actions which maximize the average reward gain in a miniblock (see methods). Critically, this *DEV* is typically close to 0 in the first few trials, i.e. there is high uncertainty on what action is the best one. In this situation, the strategy preference mostly determines the action selection of the agent. In our model, we computed the *DEV* by using forward planning, where the agent hypothetically runs simulations through all remaining future trials until the end of a miniblock, i.e. to the 15^th^ trial. The number of state space trajectories to be considered in these simulations scales exponentially with the number of remaining trials – and so does in principle the computational costs needed to simulate these trajectories. Therefore, full forward planning would be both prohibitively costly and potentially useless when the deadline is far away, rendering simpler heuristics [16] the more appropriate alternative.

It is an open question what heuristic participants actually used. In our model, the strategy preference parameter simply quantifies a preference for a parallel or sequential strategy and biases a participant’s action selection accordingly. This may mean that participants had a prior expectation whether they are going to reach G2 or just G1. Given this prior, participants could choose their action without any forward planning. In other words, to select an action in a mixed trial, participants simply assumed that they are going to reach, for example, G2. This simplifies action selection tremendously because, under the assumption that G2 will be reached, the optimal action is to use the parallel strategy at all times. To an outside observer, a participant with a strong preference for a parallel strategy may be described as overly optimistic, as this participant would choose g2-choices even if reaching G2 is not very likely, e.g. in the hard condition. Conversely, a participant with a strong preference for a sequential strategy may be described as too cautious, e.g. because that participant chooses one-goal actions in the easy condition (see S12 Fig for two example participants). Importantly, the difference in total reward between the agent and the participants is only about 5% (see Fig 3, E). This means that even though participants used a potentially suboptimal strategy preference, the impact on total reward is not that large. This is because, as we have shown, later in the miniblock, when *DEVs* become larger and are more predictive of what goal can be reached, participants choose their actions accordingly. Although we do not quantify the relative costs of full forward planning versus the observed mixture of heuristic and forward planning, we assume that an average loss of 5% of the earnings is small as compared to the reduction of computational costs when using heuristics.

There were two important features of our sequential decision making task: The first was that we used a rather long series of 15 trials to model multiple goal pursuit, where typically sequential decision making tasks would use fewer trials, e.g. 2 in the two-step task [32] with common values around 5 [21] to 8 trials [7, 8] per miniblock. The reason why we chose a rather large number of trials is that this effectively precluded the possibility that participants can plan forward and ensure that participants were exposed at least to some initial trials where they had to rely on other information than forward planning. This initial period when participants have to select actions without an accurate estimate of the future consequences of these actions is potentially most interesting for studying meta-decisions about how we use heuristics when detailed information about goal reaching probabilities is scarce. It is probably in this period of uncertainty during goal reaching, when internal beliefs and preferences have their strongest influence.

The second important feature of our task was that participants had to prioritize between two goals. This is a departure from most sequential decision making tasks, where there is typically a single goal, e.g. to collect a minimum number of points, where the alternative is a fail [7]. In our task, participants could reach one of two goals, which enables addressing questions about how participants select and pursue a specific goal, see also [9]. Our findings complement work investigating behavioural strategies for pursuing multiple goals, e.g. [33], showing that pursuit strategies depend on environmental characteristics, subjective preferences and changes in context when getting closer to the goal. In line with our findings, a recent study [34] showed that decisions whether to redress the imbalance between two assets or to focus on a distinct asset during sequential goal pursuit were best fit by a dynamic programming model with a limited time horizon of 7.5 trials (20 trials would be the optimum). In future research, the pursuit of multiple goals in sequential decision making tasks may also be a basis for addressing questions about cognitive control during goal-reaching, e.g. how participants regulate the balance between stable maintenance and flexible updating of goal representations [35].

Another important factor when modelling the use of forward planning is that complexity and time can, in principle be dissociated. For example, a temporally distant goal might have only low planning complexity because one must consider only a few decision sequences leading to the goal. Conversely, a temporally proximate goal might have high planning complexity because of a large number of potential actions sequences that may lead to the goal. In future research, by testing sequential tasks with varying branching factor (number of potential actions in each trial) one could selectively test how time to goal and planning complexity influence the arbitration of forward planning and the use of heuristics.

It is unclear what mechanism made participants actually use a strategy preference different from zero in our task. It is tempting to assume that participants might have used their usual approach, which they might apply in similar real-life situations, to select their goal strategies when the computational costs of forward planning are high and the prediction accuracy is low. In other words, participants who had a preference for a parallel strategy might either show a tendency towards working on multiple goals at the same time or entertain the belief that tasks should be approached with an optimistic stance. Conversely, participants with a preference for a sequential strategy might have made good experiences with using a more cautious approach and would tend to pursue one goal after the other.

We would like to note that the proposed model does not explicitly model the arbitration between forward planning and heuristic decision making. The computational model to fit participant behaviour uses at its core full forward planning as the optimal agent does. The effect of strategy preference just changes the action selection result, but the underlying computation to determine the *DEV* is still based on forward planning. Clearly, if a real agent used our model, this agent would not save any computations because forward planning is still used for all trials. The open question is how an agent makes a meta-decision to not use goal-directed forward planning but to rely on heuristics and other cost-efficient action selection procedures [11]. To make this meta-decision, an agent cannot rely on the *DEV* because this value is computed by forward planning. An alternative way would be to use an agent’s prior experience to decide that the goal is still too temporally distant to make an informed decision with an acceptable computational cost. Such a meta-decision would depend on several factors, e.g. the relevance of reaching G2, intrinsic capability and motivation of planning forward, or a temporal distance parameter which signals urgency to start planning forward. In the future we plan to develop such meta-decision-making models and predict the moment at which forward planning takes over the action selection process.

It is also possible that participants use, apart from simple heuristics, other approximate planning strategies to reduce computational costs. For example, one could sample only a subset of sequences to compute value estimates. Indeed, in another study it was found that participants prune a part of the decision tree in response to potential losses, even if this pruning was suboptimal [36]. Another important point is that the planning process itself might be error-prone and therefore value calculations over longer temporal horizons may be noisier. This could presumably account for temporal modulations of the precision parameter β. In future work one could test for evidence of alternative planning algorithms that allow to sample subsets of (noisy) forward planning trajectories to further delineate how humans deal with computational complexity in goal-directed decision scenarios.

Taken together, the present research shows that over prolonged goal-reaching periods, individuals tend to behave in a way that approaches the behaviour of an optimal agent, with noticeable differences early in the goal-reaching period, but nearly optimal behaviour when the goal is close. It also highlights the potential of computational modelling to infer the decision parameters individuals use during different stages of sequential decision-making. Such models may be a promising means to further elucidate the dynamics of decision-making in the pursuit of both laboratory and everyday life goals.

## Supporting information

S1 Movie. Simulated goal success and total reward for variable precision parameter

S2 Movie. Simulated suboptimal g-choices for variable precision parameter

S3 Movie. Simulated goal success and total reward for variable strategy preference parameter

S4 Movie. Simulated suboptimal g-choices for variable strategy preference parameter

S1 Notebook. Parameter recovery simulations

## Supporting information

**S1 Table.** Classification of accept-wait responses into either two-goal-choices (g2) or one-goal-choices (g1).

| Offer | Points | Response | Classification |
| --- | --- | --- | --- |
| Ab | $Pts_t^A - Pts_t^B > 1$ | accept | g1 |
| Ab | $Pts_t^A - Pts_t^B > 1$ | wait | g2 |
| Ab | $Pts_t^A - Pts_t^B < -1$ | accept | g2 |
| Ab | $Pts_t^A - Pts_t^B < -1$ | wait | g1 |
| Ab | $Pts_t^A - Pts_t^B = 1$ | accept | g1 |
| Ab | $Pts_t^A - Pts_t^B = 1$ | wait | g2 |
| Ab | $Pts_t^A - Pts_t^B = -1$ | accept | nan |
| Ab | $Pts_t^A - Pts_t^B = -1$ | wait | nan |
| Ab | $Pts_t^A - Pts_t^B = 0$ | accept | g1 |
| Ab | $Pts_t^A - Pts_t^B = 0$ | wait | g2 |
| aB | $Pts_t^A - Pts_t^B > 1$ | accept | g2 |
| aB | $Pts_t^A - Pts_t^B > 1$ | wait | g1 |
| aB | $Pts_t^A - Pts_t^B < -1$ | accept | g1 |
| aB | $Pts_t^A - Pts_t^B < -1$ | wait | g2 |
| aB | $Pts_t^A - Pts_t^B = 1$ | accept | nan |
| aB | $Pts_t^A - Pts_t^B = 1$ | wait | nan |
| aB | $Pts_t^A - Pts_t^B = -1$ | accept | g1 |
| aB | $Pts_t^A - Pts_t^B = -1$ | wait | g2 |
| aB | $Pts_t^A - Pts_t^B = 0$ | accept | g1 |
| aB | $Pts_t^A - Pts_t^B = 0$ | wait | g2 |

**S1 Fig.**
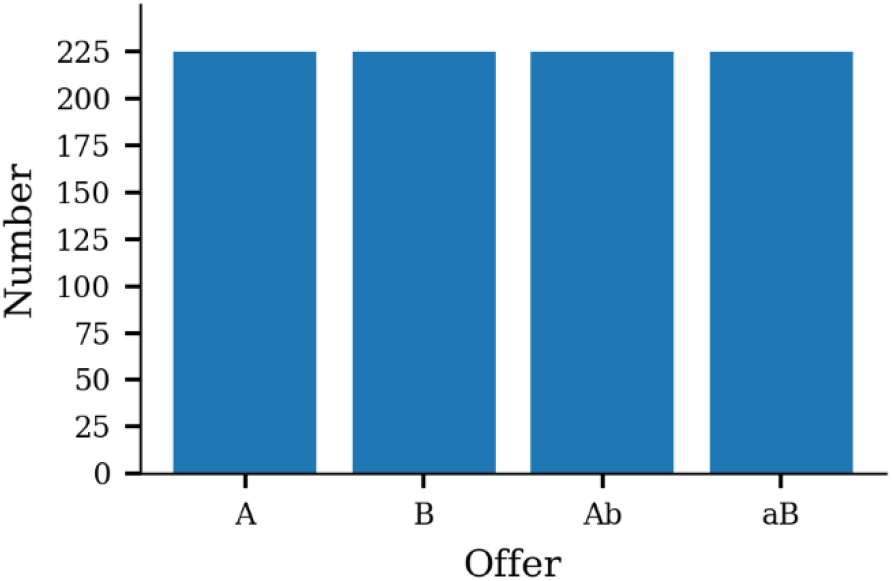
Occurrence of offer types across all 900 trials.

**S2 Fig.**
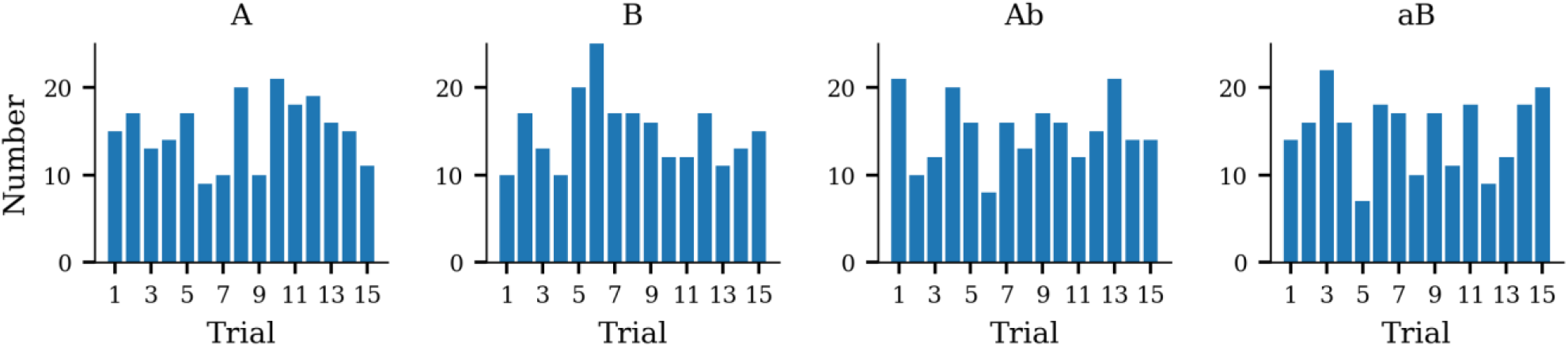
Occurrence of offer types binned with respect to trial.

**S3 Fig.**
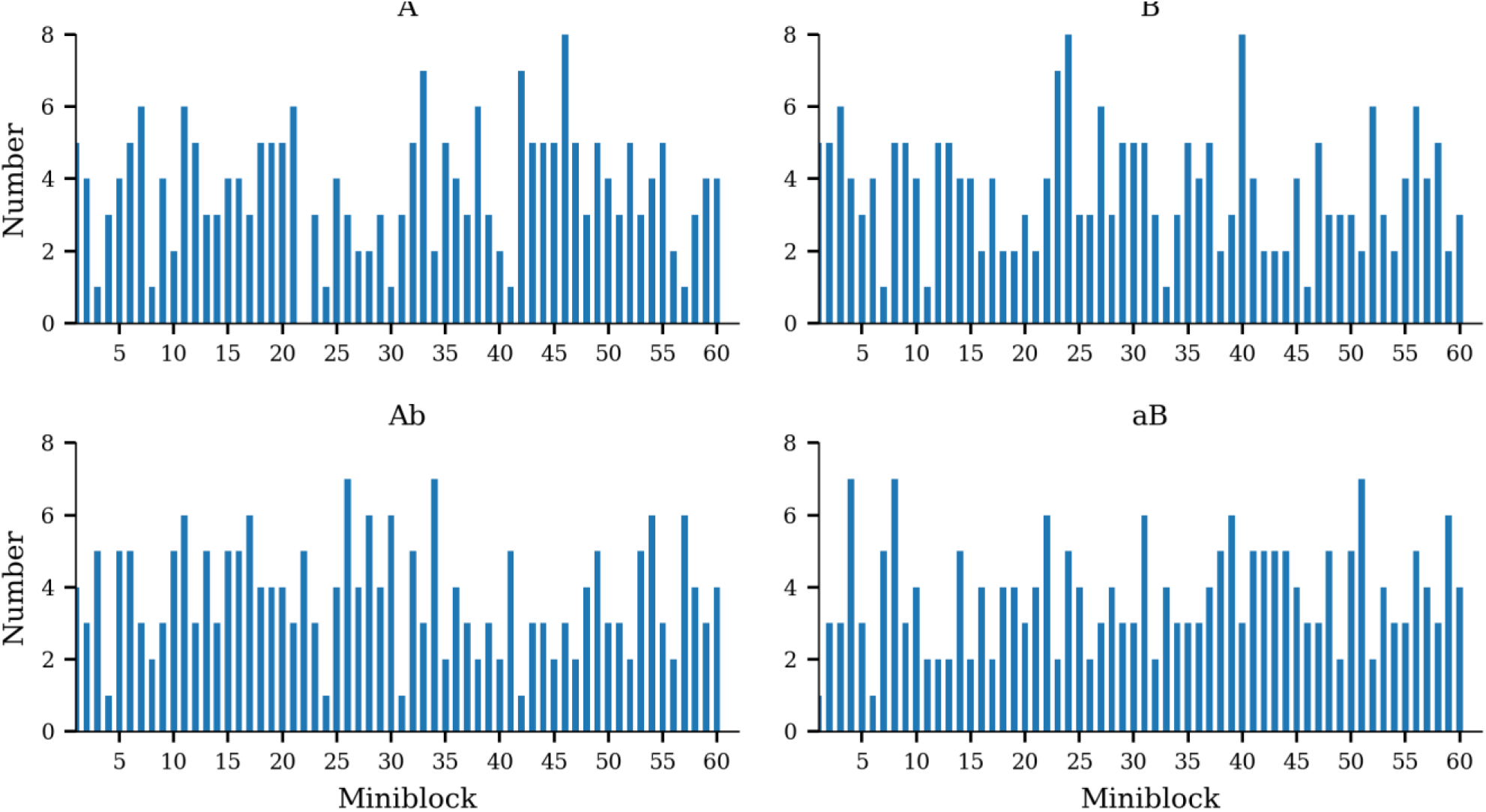
Occurrence of offer types binned with respect to miniblock.

**S4 Fig.**
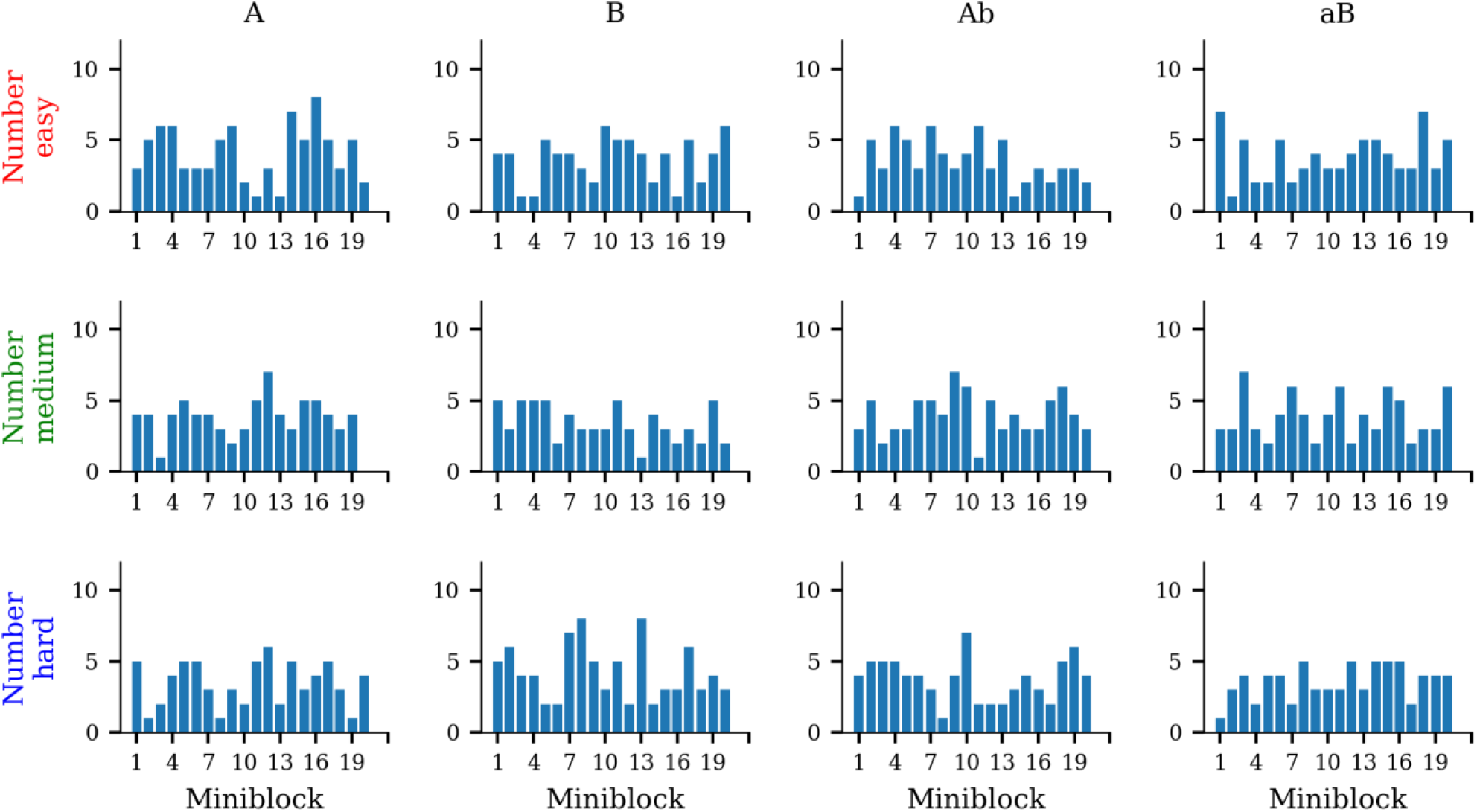
Occurrence of offer types binned with respect to miniblock and difficulty.

**S5 Fig.**
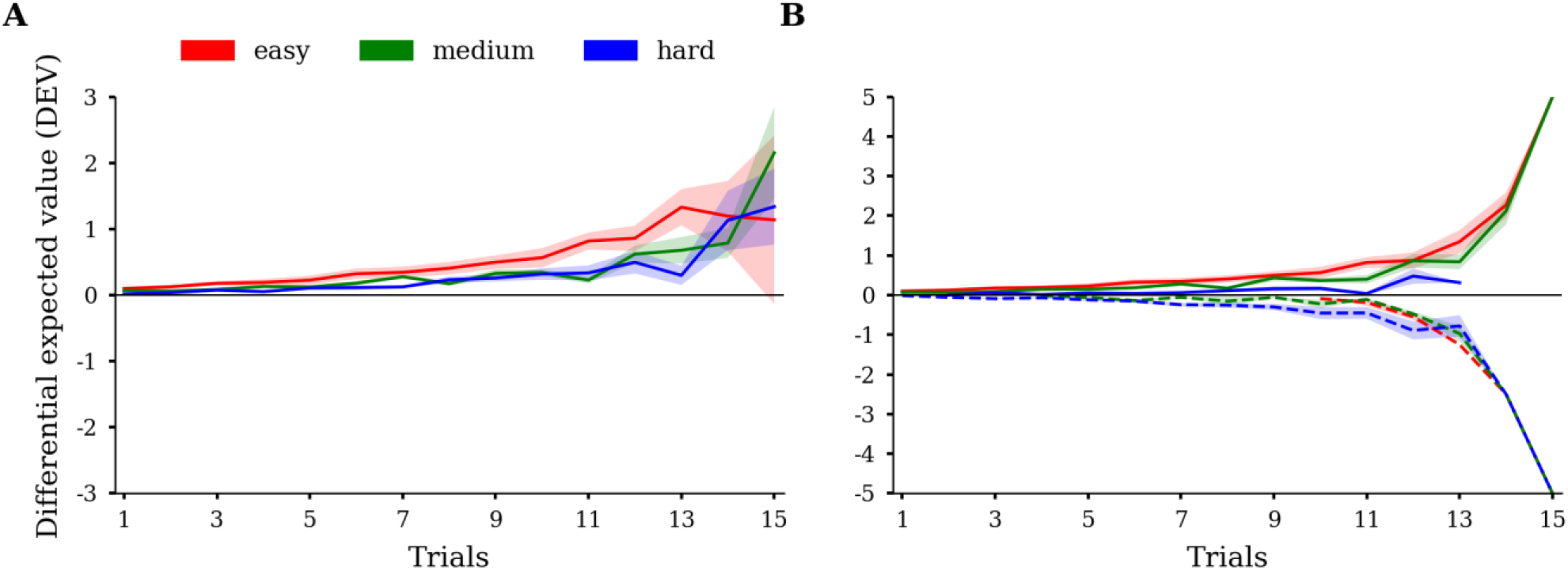
Average absolute (A) and signed (B) differential expected value (*DEV*) per trial and condition. Discount and reward ratio had been fixed (*γ* = 1, *κ* = 1). Average absolute *DEVs* at the beginning of the miniblock are smaller than in the end, indicating the relative importance of decisions close to the final trial of miniblocks. Conditions are colour coded. The shaded areas represent SD.

**S6 Fig.**
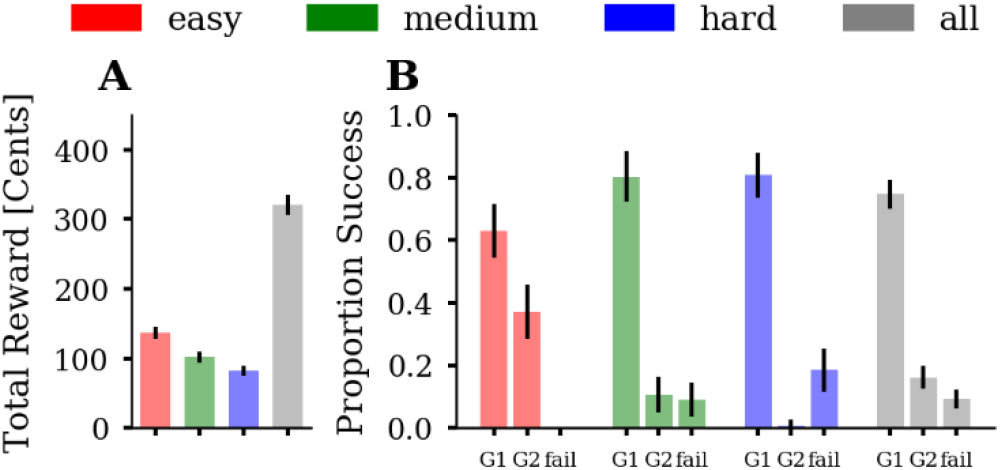
Simulated goal success and total reward of a random agent that always accepts basic offers but guesses for mixed offers (*θ* = 0, *β* → 0, *γ* = 1, *κ* = 1). **(A)** Average total reward across agent instances (n =1000). **(B)** Proportion of successful goal-reaching, averaged across agent instances, for each of the three conditions. We plot the proportion of reaching, at the end of a miniblock, a single goal (G1), both goals (G2), or no goal (fail). The random agent achieves fewer G2-successes in easy and medium than the participants but fails more often in medium and hard. The three conditions are colour-coded (easy = red, medium = green, blue = hard) and the average over conditions is shown in grey. Error bars depict SD.

**S7 Fig.**
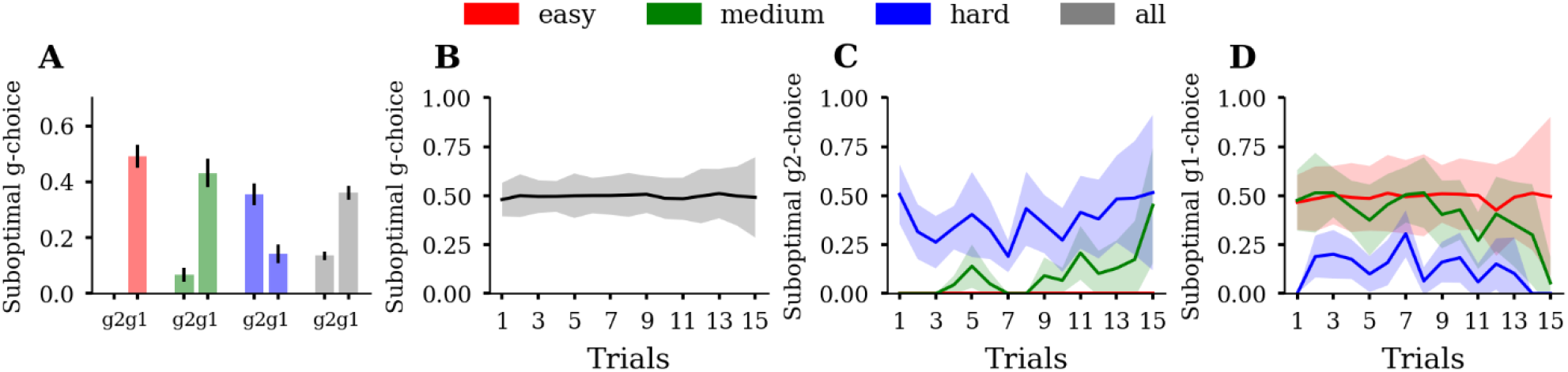
Simulated suboptimal g-choices of a random agent that always accepts basic offers but guesses for mixed offers (*θ* = 0, *β* → 0, *γ* = 1, *κ* = 1). **(A)** Proportions of suboptimal g1-choices (g1) and suboptimal g2-choices (g2), averaged over agent instances (n =1000). The random agent makes many suboptimal g1-choices in the easy and medium and many suboptimal g2-choices in the hard conditions. Summing together g1 and g2 yields approximately 50% suboptimal g-choices. **(B)** Suboptimal g-choices as a function of trial averaged over agent instances. The random agent makes approximately 50% suboptimal g-choices across all trials in the miniblock. If participants use non-random response strategies, i.e. planning or heuristics, their pattern of suboptimality across trials should deviate from the straight-line pattern of the random agent. **(C)** Suboptimal g2-choices as a function of trial averaged over agent instances. **(D)** Suboptimal g1-choices as a function of trial averaged over agent instances. Summing together g1 (D) and g2 (C) yields approximately 50% suboptimal g-choices across trials. Error bars and shaded areas depict SD. Conditions are colour coded.

**S8 Fig.**
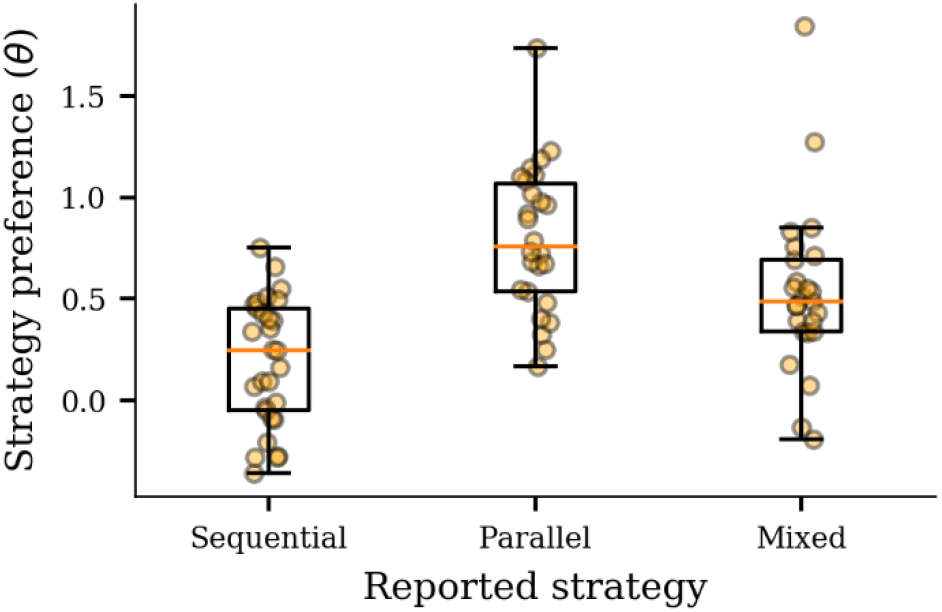
Qualitative comparison of participants’ reported strategy use and fitted strategy preference parameter. Participants who reported the use of a sequential strategy had lower estimated strategy preference, including the most negative values, than participants who reported the use of a parallel strategy. Participants who reported mixed use of a parallel and sequential strategy had greater strategy preference than the sequential group but lower estimates than the parallel group. The plot shows 80 of 89 participants whose verbal reports matched with one of the three strategy categories.

**S9 Fig.**
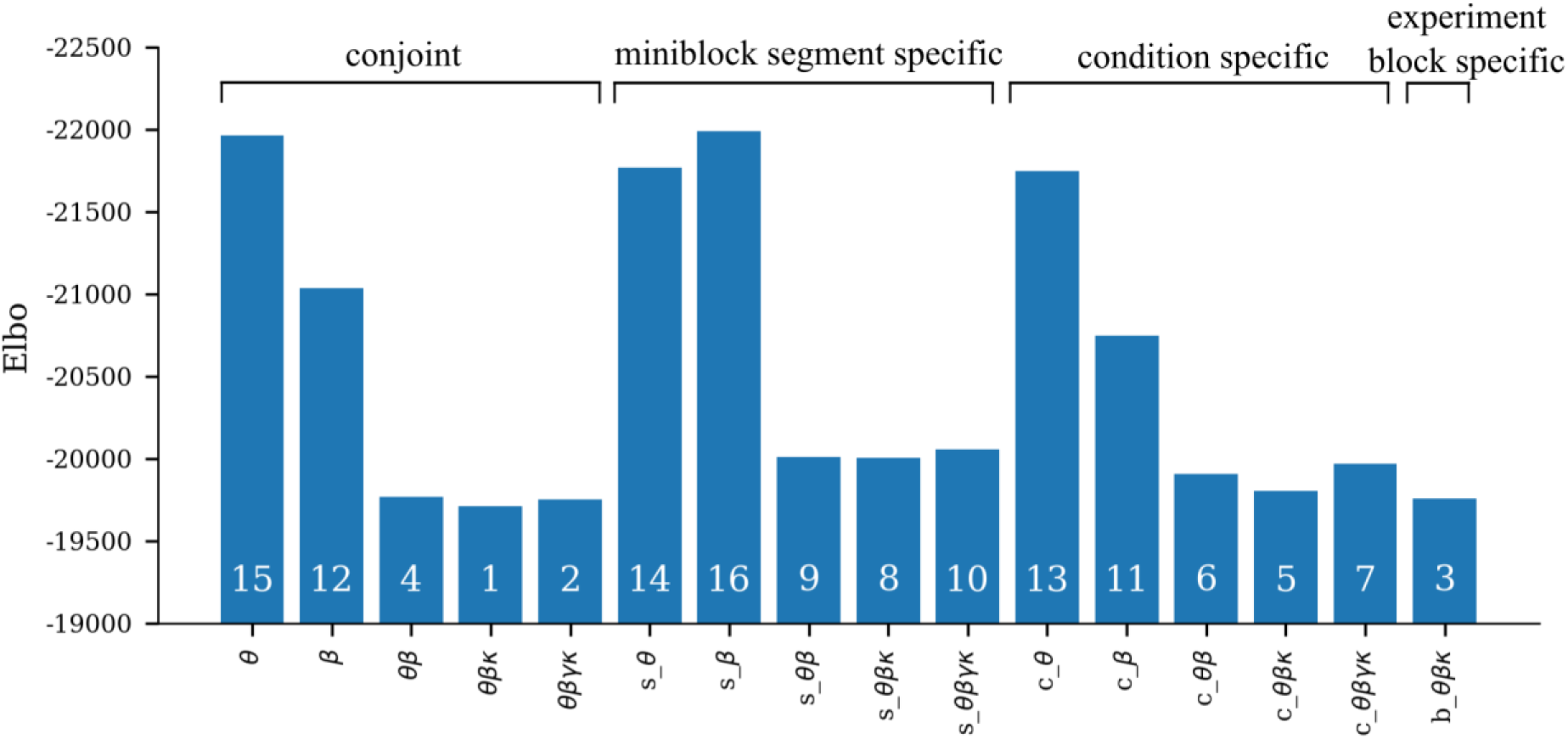
Comparing Elbo (evidence lower bound) between different model variants. White numbers represent the rank from highest to lowest Elbo. Model comparisons showed that the three parameter model (*θ, β, κ*) had the highest model evidence. Adding *γ* did not increase model evidence (*elbo_θβκ_* – *elbo_θβγκ_* = −44). Estimating model parameters separately for miniblock segments (trial 1-5, trial 6-10, trial 11-15; prefix ‘s_’ in the figure) had lower model evidence compared to the winning model (*elbo_θβκ_* – *elbo_s_θβκ_* = −294). Estimating model parameters separately for conditions (easy, medium, hard; prefix ‘c’ in the figure) had lower model evidence compared to the winning model (*elbo_θβκ_* – *elbo_c_θβκ_* = −94). Estimating model parameters separately for experiment blocks (miniblock 1-20, miniblock 21-40, miniblock 41-60; prefix ‘b’ in the figure) had also lower model evidence compared to the winning model (*elbo_θβκ_* – *elbo_s_θβκ_* = −48). Bars in the plot depict Elbo averaged over the last 20 posterior samples.

**S10 Fig.**
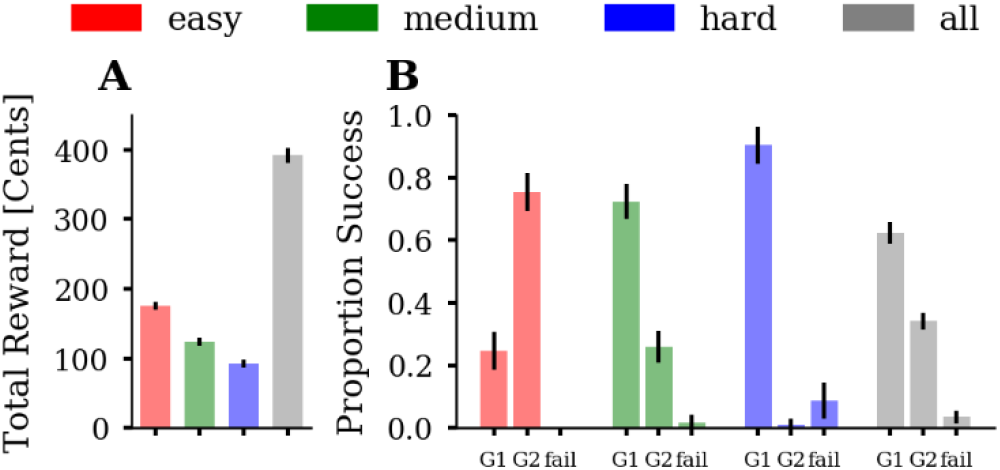
Posterior predictive checks: Simulated goal success and total reward closely resemble observed participant behaviour. **(A)** Average total reward across samples (n = 1,000). **(B)** Proportion of successful goal-reaching, averaged across samples, for each of the three conditions. We plot the proportion of reaching, at the end of a miniblock, a single goal (G1), both goals (G2), or no goal (fail). The three conditions are colour-coded (easy = red, medium = green, blue = hard) and the average over conditions is shown in grey. Error bars depict SD. Data were generated using 1,000 posterior samples from the group hyper parameters.

**S11 Fig.**
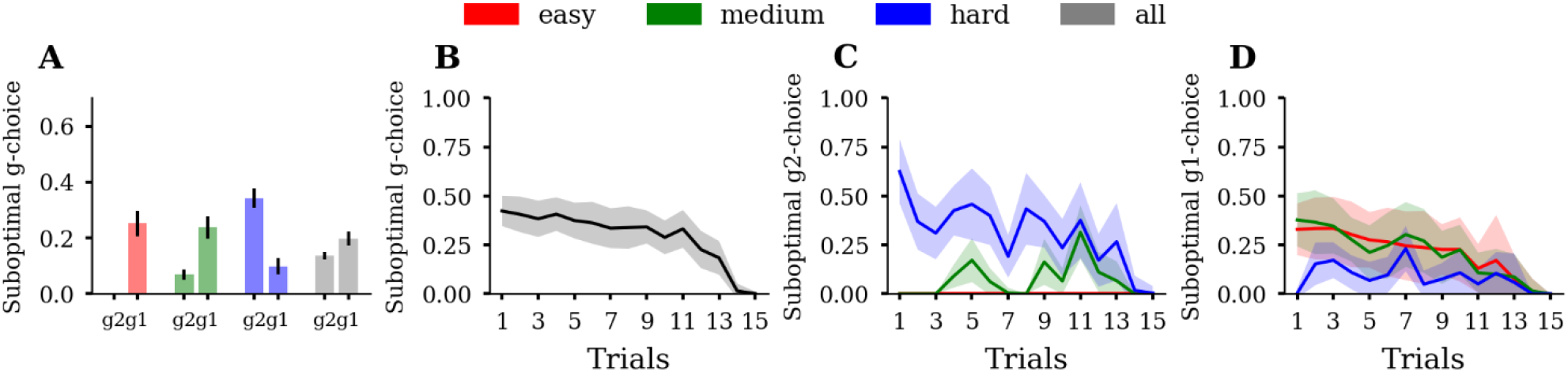
Posterior predictive checks: Simulated suboptimal g-choices closely resemble observed participant behaviour. **(A)** Proportions of suboptimal g1-choices (g1) and suboptimal g2-choices (g2), averaged over samples (n =1,000). **(B)** Suboptimal g-choices as a function of trial averaged over samples. **(C)** Suboptimal g2-choices as a function of trial averaged over samples. **(D)** Suboptimal g1-choices as a function of trial averaged over samples. Error bars and shaded areas depict SD. Conditions are colour coded. Data were generated using 1,000 posterior samples from the group hyper parameters.

**S12 Fig.**
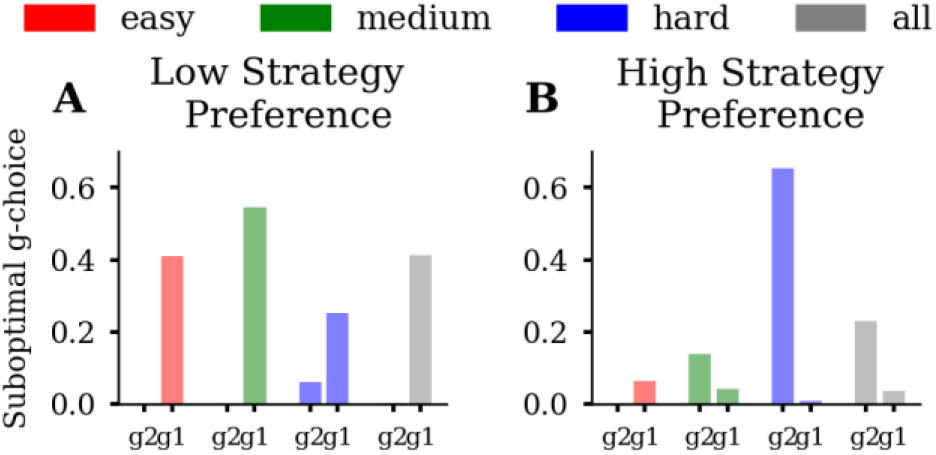
Comparison of suboptimal g-choices between a low strategy preference and high strategy preference participant. The plot shows proportions of suboptimal g1-choices (g1) and suboptimal g2-choices (g2) **(A)** of the participant with the lowest fitted strategy preference (*θ* = −0.36) and **(B)** of the participant with the highest fitted strategy preference (*θ* = 1.84). The low strategy preference participant prefers a sequential strategy leading to suboptimal g1-choices in the easy and medium condition. The participant with a high strategy preference parameter prefers a parallel strategy, resulting in a few suboptimal g1-choices in easy in and medium but a large number of suboptimal g2-choices in the hard condition.

## S1 Text: Task instructions (translated from German)

Dear participant, your task in this experiment is to reach goals. Within a block, consisting of 15 trials, you can either reach goal A, goal B or both goals at the same time. For one reached goal you will gain additional 5 Cents and for two reached goals additional 10 Cents. Your task is to obtain as much money as possible.

To reach goals, you must collect points. You can get points by accepting an offer. Some offers however, might have a negative effect on the state of a goal. Your task is to decide in every trial, whether to accept an offer or wait for the next offer. Press “up arrow” to accept an offer and “down arrow” to wait.

### Important

Please decide deliberately but speedily. If you decide too slowly, you will get a notification. After every 5 notifications, 50 Cents will be subtracted from your bonus-payout. (The experiment starts with a training phase, in which no money can be lost.)

### More about the goals

Your goal progress will be represented by a bar, which is labelled with A or B. A goal counts as achieved, if one of the bars reaches or surpasses the white horizontal mark. The goal state will be evaluated after the end of the 15 trials.

### More about the offers

There are 4 different offers – A, B, Ab an aB. All offers have the same occurrence probability of 25%. The offers differ with respect to their effect on the goal state. A increases the A-bar by one point. B increases the B-bar by one point. Ab increases the A-bar by one point and subtracts one point from the B-bar. aB increases the B-bar by one point and subtracts 1 point from the A-bar.

### Initial conditions

At the beginning of the block, you already have some A- and B-points. The amount of initial points varies from block to block.

**S1 Movie. Simulated goal success and total reward where the precision parameter *β* varies between 0.25 and 3 with *θ, γ*, and *κ* sampled from their fitted population mean. (A)** Average total reward across agent instances (n =1,000). An increase in *β* increases total reward obtained in the easy and medium but decreases total reward in the hard condition. **(B)** Proportion of successful goal-reaching, averaged across agent instances, for each of the three conditions. We plot the proportion of reaching, at the end of a miniblock, a single goal (G1), both goals (G2), or no goal (fail). An increase in *β* increases G2 success rate in easy and medium but also increases fail rate in medium and hard. The three conditions are colour-coded (easy = red, medium = green, blue = hard) and the average over conditions is shown in grey. Error bars depict SD.

**S2 Movie. Simulated suboptimal g-choices where the precision parameter *β* varies between 0.25 and 3 with *θ, γ*, and *κ* sampled from their fitted population mean. (A)** Proportions of suboptimal g1-choices (g1) and suboptimal g2-choices (g2), averaged over agent instances (n =1000). An increase in *β* decreases suboptimal g1- and g2-choices. **(B)** Suboptimal g-choices as a function of trial averaged over agent instances. The influence of *β* and the associated decrease of suboptimal g-choices successively increases towards the end of the miniblock. Suboptimal g-choices in the first half of the miniblock are largely unaffected by the *β* parameter. **(C)** Suboptimal g2-choices as a function of trial averaged over agent instances. An increase in *β* decreases suboptimal g2-choices late in the miniblock in medium and hard but not in easy. **(D)** Suboptimal g1-choices as a function of trial averaged over agent instances. An increase in *β* decreases suboptimal g1-choices late in the miniblock in easy and medium but not in hard. Error bars and shaded areas depict SD. Conditions are colour coded.

**S3 Movie. Simulated goal success and total reward where the strategy preference parameter *θ* varies between −1 and 1 with *β, γ*, and *κ* sampled from their fitted population mean. (A)** Average total reward across agent instances (n =1000). An increase in *θ* increases total reward obtained in easy and medium but decreases total reward in hard. **(B)** Proportion of successful goal-reaching, averaged across agent instances, for each of the three conditions. We plot the proportion of reaching, at the end of a miniblock, a single goal (G1), both goals (G2), or no goal (fail). An increase in *θ* increases G2 success rate in easy and medium but also increases fail rate in medium and hard. The three conditions are colour-coded (easy = red, medium = green, blue = hard) and the average over conditions is shown in grey. Error bars depict SD.

**S4 Movie. Simulated suboptimal g-choices where the strategy preference parameter *θ* varies between −1 and 1 with *β, γ*, and *κ* sampled from their fitted population mean. (A)** Proportions of suboptimal g1-choices (g1) and suboptimal g2-choices (g2), averaged over agent instances (n =1000). An increase in *θ* decreases suboptimal g1-choices and increases suboptimal g2-choices. Suboptimal g1-choices decrease more in easy and medium than in hard. Suboptimal g2-choices decrease more in hard than in easy and medium. **(B)** Suboptimal g-choices as a function of trial averaged over agent instances. A change in *θ* affects the number of suboptimal g-choices made at the beginning but not at the end of the miniblock. For *θ* > 0 suboptimal g-choices further decrease, because g2-choices are often optimal in easy and medium. **(C)** Suboptimal g2-choices as a function of trial averaged over agent instances. An increase in *θ* increases suboptimal g2-choices early in the miniblock, predominantly in the hard condition. **(D)** Suboptimal g1-choices as a function of trial averaged over agent instances. An increase in *θ* decreases suboptimal g1-choices early in the miniblock, predominately in easy and medium. Error bars and shaded areas depict SD. Conditions are colour coded.

**S1 Notebook. Parameter recovery simulations.**

