## Supplementary material for "Dynamic integration of forward planning and heuristic preferences during multiple goal pursuit": S1 Notebook. Parameter recovery simulations

### Testing inference

November 6, 2019

Here we will look into the inference of different hierarchical parametric models one could apply to the backward induction agent used for the two-goal task.

The free parameters of the model are as follows:

- beta (precision) $\beta$
- bias (strategy preference) $\theta$
- discount rate $\gamma$
- reward ratio $\kappa$

Note that we assumed that agent always accepts A, B offer.

We are interested in comparing different parametric models in their accuracy, that is, ability to accurately recover model parameters. For this purpose we will vary only prior over model parameters $p(\Phi)$, where $\Phi$ denotes a set of model parameters and their hyper priors. Furthermore, we will perform inference in the unconstrained parameter space.

We will first simulate some behavioral data for $n = 89$ subjects, where we define for each subject a different set of free model parameters, obtained from the means of the posterior estimates based on the empirical data.

[1]:
```
import torch
import pyro
pyro.validation_enabled(True)

import numpy as np
import seaborn as sns
import matplotlib.pyplot as plt
import pandas as pd

from tasks import TwoGoalTask
from agents import Informed
from simulate import Simulator
from inference import Inferrer

from helpers import offer_state_mapping

zeros = torch.zeros
ones = torch.ones
```

[2]:
```python
# load posterior estimates
sub_sample = pd.read_csv('../Results/theta_beta_gamma_kappa/
↪subject_posterior_sample_20191022-141145.csv',index_col=0)
x_sample = sub_sample.groupby('subject').mean()
sns.catplot(data=x_sample, aspect=2);
```

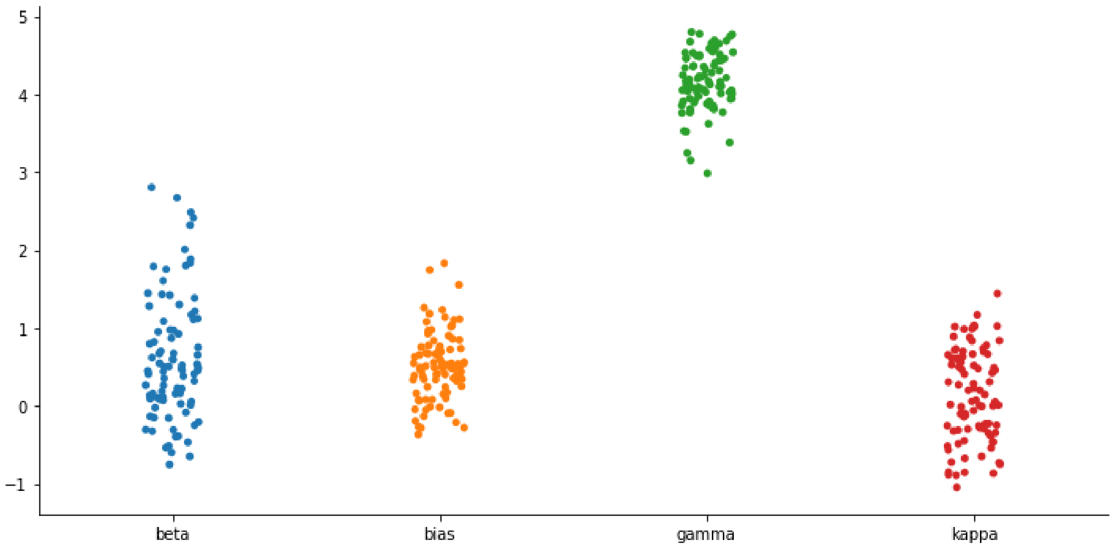


[ ]:
```python
torch.manual_seed(1234)  # fix seed for reproducability

runs = x_sample.shape[0]  # number of subjects
mini_blocks = 60  # number of mini-blocks
trials = 15  # number of trials per mini block
na = 2  # number of choices in each trial
no = 4  # number of different offers (A, B, Ab, Ba)
threshold = 10  # goal threshold
lam = 5  # reward strength of the single goal
offer_prob = torch.FloatTensor([1/no, 1/no, 1/no, 1/no])  # offer probability

# Get initial states
tmp = pd.read_csv('../Results/preprocessed_results.csv')
tmp = tmp.loc[tmp['phase'] > 1]
trial_values = tmp.loc[:, 'score_A_before':'score_B_before'].values - 1

offers = (tmp['offer'].values - 1).reshape(89, -1, trials)[0]
offers = torch.tensor(offers).repeat(runs, 1, 1)

L = trial_values.max()+1
ns = np.asscalar(L**2) #number of states
```

```
states = torch.tensor(np.ravel_multi_index(trial_values.T, (L, )*2).T.
 ↪reshape(89, -1, trials))
initial_states = torch.zeros(runs, mini_blocks, dtype=torch.long)
initial_states[:] = states[0, :, 0][None, :]

#define state transition matrix for the environment p(s'|s, o, a)
offer_state_tm = zeros(na, no, ns, ns)
#note that the order of elements is inverted -> a, o, s| s'

#a = 0 corresponds to wait action, states don't change in this case
offer_state_tm[0] = torch.eye(ns).repeat(no, 1, 1)

#a = 1 coresponds to accepting an offer
offer_state_tm[1] = offer_state_mapping(L, no)

#set state reward mapping on the end trial R(s)
outcomes = np.sum(np.indices((L, L)) >= threshold, axis=0).flatten()
outcomes = outcomes*lam
outcomes = torch.from_numpy(outcomes).float()

env = TwoGoalTask(offer_state_tm,
                  runs=runs,
                  mini_blocks=mini_blocks,
                  trials=trials,
                  number_states=ns,
                  number_offers=no,
                  initial_states=initial_states,
                  offers=offers)

x = torch.zeros(runs, 6)

# fix parameters associated with A,B offer types. Agent always accepts A,B␣
 ↪offers.
x[:, 0] = -100.
x[:, 2] = 100.

# asign parameter values in the transformed parameter space
x[:, 1] = torch.from_numpy(x_sample.beta.values)
x[:, 3] = torch.from_numpy(x_sample.bias.values)
x[:, 4] = torch.from_numpy(x_sample.gamma.values)
x[:, 5] = torch.from_numpy(x_sample.kappa.values)

agent = Informed(offer_state_tm,
                 outcomes,
                 offer_prob,
                 runs=runs,
                 mini_blocks=mini_blocks,
```

```
                 trials=trials,
                 na=na,
                 ns=ns,
                 no=no).set_parameters(x)

sim = Simulator(env, agent, runs=runs, mini_blocks=mini_blocks, trials=trials)
sim.simulate_experiment()

stimulus = {'states':env.states, 'offers': env.offers}
responses = sim.responses

# set valid states to all except G2 states (the ones in which one gets 10␣
 ↪points)
valid = ~(outcomes[env.states[..., :-1]] == 10)

map_states_to_points = torch.tensor(
        np.array(np.unravel_index(np.arange(ns), (L, L))).T
        )
points = map_states_to_points[env.states[..., :-1]]

point_diff = points[..., 0] - points[..., 1]

exclude = ((point_diff == 1) & (env.offers == 3)) | ((point_diff == -1) & (env.
 ↪offers == 2))

mask = valid & ~exclude
mask *= env.offers > 1

agent = Informed(offer_state_tm,
                 outcomes,
                 offer_prob,
                 runs=runs,
                 mini_blocks=mini_blocks,
                 trials=trials,
                 na=na,
                 ns=ns,
                 no=no).set_parameters(x)

# fixed beta_1 and theta_1 parameters
vals = torch.ones(runs, 2)
vals[:, 0] = -100.
vals[:, 1] = 100.
fixed_params = {'labels': [0, 2],
                'values': vals}
```

First we will test a non-hierarchical model, where we use a Gaussian prior over free-model parameters (transformed to an unconstrained space). If we denote with $\vec{x} = (x_1, \ldots, x_d)$ vector of free model parameters, then the prior over all subjects $p(\hat{x})$ is defined

$$p(\hat{x}) = \prod_{n=1}^{N} \prod_{i=1}^{d} \mathcal{N}\left(x_i^n; m_i, s_i\right)$$

where $d$ denotes number of free model parameters and $n$ subject whose set of responses we are fitting to the model.

In the case of the flat prior the approximate posterior factorises as follows

$$p(\hat{x}|R, O) \approx Q(\hat{x}|\theta) = \prod_{n=1}^{N} \mathcal{N}_d(\vec{\mu}^n, \Sigma^n)$$

where $d$ denotes the number of free parameters.

[4]:
```
infer_nh = Inferrer(agent, stimulus, responses, mask=mask,␣
 ↪fixed_params=fixed_params)
infer_nh.infer_posterior(iter_steps=200, parametrisation='non-hierarchical')
```

```
Mean ELBO 19340.12: 100%|      | 200/200 [35:57<00:00, 10.86s/it]
```

[5]:
```
#plot posterior estimates in comparison with true values
labels = ['beta', 'bias', 'gamma', 'kappa']
res = infer_nh.sample_from_posterior(labels, n_samples=999)

trans_pars_nh = res[0].melt(id_vars='subject', var_name='parameter')
loss_nh = infer_nh.loss

#plot convergence of stochasitc ELBO estimates (log-model evidence)
plt.figure()
plt.plot(loss_nh[-150:])

fig, axes = plt.subplots(1, len(labels), figsize=(15, 5))
m = res[0].groupby('subject').mean()
std = res[0].groupby('subject').std()

for i,l in enumerate(labels):
    axes[i].errorbar(x_sample[l], m[l], std[l], c='r', marker='o', linestyle='')
    axes[i].plot(x_sample[l], x_sample[l], 'k--')
    axes[i].set_title(l)
    axes[i].set_xlabel('true value')

axes[0].set_ylabel('estimated value')
```

[5]:
```
Text(0, 0.5, 'estimated value')
```

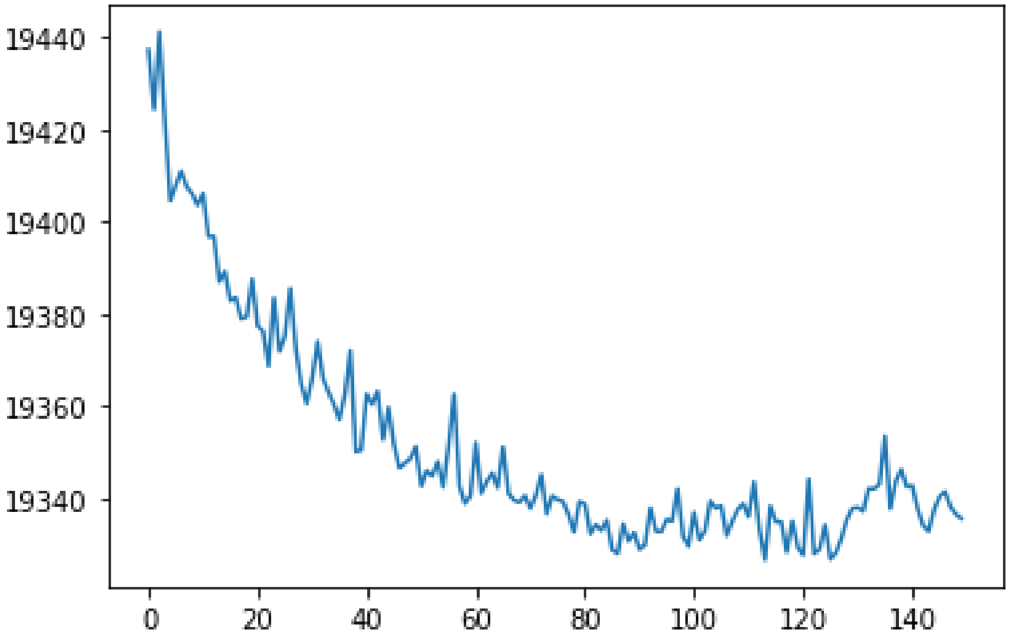


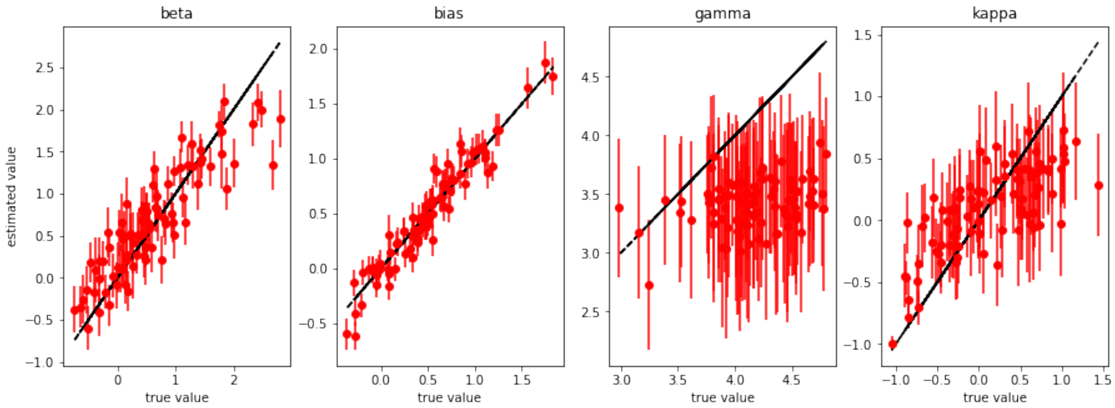


[6]:
```
# Estimating probability that a true value falls within 95% probability␣
↪interval.
def posterior_accuracy(labels, df, z):
    for i, lbl in enumerate(labels):
        std = df.loc[df['parameter'] == lbl].groupby(by='subject').std()
        mean = df.loc[df['parameter'] == lbl].groupby(by='subject').mean()
        print(lbl, np.sum(((mean+2*std).values[:, 0] > z[i])*((mean-2*std).
↪values[:, 0] < z[i]))/runs)
```

[7]:
```
names = ['beta', 'bias', 'gamma', 'kappa']
true_vals = x_sample.values.T
```

```
posterior_accuracy(names, trans_pars_nh, true_vals)

# compute probability that a sample from posterior is smaller than the
# true value and plot the distribution summed over subjects.
# In ideal scenario we should get a uniform distribution for all parameters
fig, axes = plt.subplots(1, 4, figsize=(15, 5))
for i, l in enumerate(labels):
    R = res[0][l].values.reshape(-1, runs) < true_vals[i]
    axes[i].hist(R.sum(0), bins=20)
    axes[i].set_title(l)
```

```
beta 0.8651685393258427
bias 0.9550561797752809
gamma 0.9101123595505618
kappa 0.9213483146067416
```

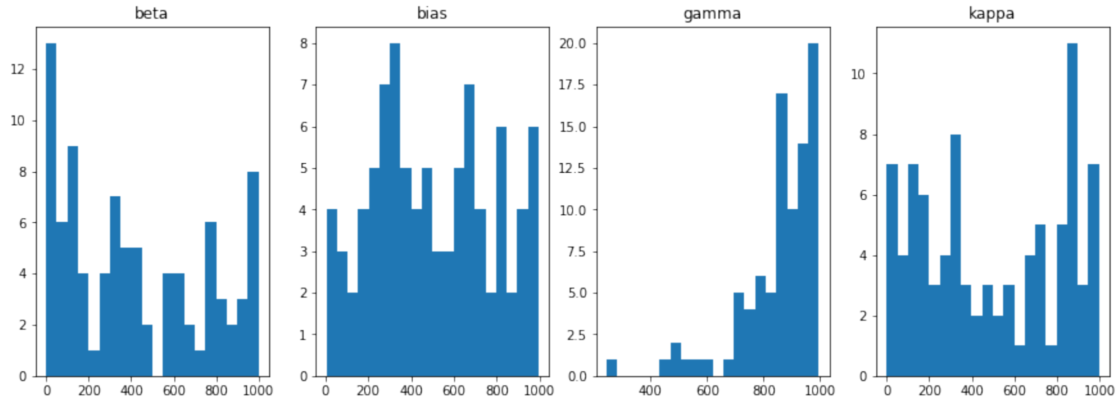


Note that in the above plot we have a good estimate of the biases, but very noisy estimates of other parameters after 200 iterations. In practice, one would expect that a non-hierarchical model has more noiser estimates and potential exhibits slower convergance towards the free energy minimum, compared to hierarchical parametric models.

Here, we will define and test a hierarchical parametric model, where we will use the so-called horseshoe prior as the prior of hyper-parameters. The horseshoe prior assumes that each parameter might have different prior uncertainty, which is independent of the specific subject (all subjects share the same prior uncertainty for a specific parameter). In addition, we will also assume that each parameter is sampled from a different prior mean, where we also assume that the prior mean is defined on the group level, hence

$$\mu_i \sim \mathcal{N}(m_i, s_i)$$
$$\sigma_i \sim C^+(0, 1)$$
$$x_i^n \sim \mathcal{N}(\mu_i, \lambda\sigma_i)$$

where $i \in \{1, \ldots, d\}$ and $n \in \{1, \ldots, N\}$, $C^+(0,1)$ denotes a Half-Cauchy distribution with scale $b = 1$. Note that parameters $m_i$, $s_i$, and $\lambda$ become new hyper parameters of the hierarchical model which are also estimated from the data, using the empirical Bayes procedure. In other words, we are performing a points estimate of $\eta = (m_1, \ldots, m_d, s_1, \ldots, s_d, \lambda)$ in parallel to the posterior estimate of $\vec{x}^n$, $\vec{\mu}$, and $\vec{\sigma}$. Formally, this becomes equivalent to doing a maximal likelihood estimate over marginal likelihood

$$\ln p(D|\eta^*) = \arg\max_{\eta} \ln p(D|\eta) \approx \arg\max_{\eta,\gamma} ELBO(\eta, \gamma)$$

where $ELBO(\eta, \gamma)$ denotes negative variational free energy (evidence lower bound) expressed as

$$ELBO = E_Q \left[ \ln \frac{p(D, Z|\eta)}{Q(Z|\gamma)} \right]$$

To define the posterior distribution over model parameters and hyper parameters, we follow the fact that the the true joint posterior contains dependencies between free parameters. To account for this we will approximate the posterior in the form of the multivariate normal distribution, which is factorised over global hyper parameters, and local parameters for each subject. Hence,

$$Q(Z|\gamma) = Q(\vec{\mu}, \vec{\sigma}|\gamma') \prod_n Q(\vec{x}_n|\gamma''_n)$$

We express the approximate posterior over group parameters $(\vec{\mu}, \vec{\sigma})$ as

$$Q(\vec{\mu}, \vec{\sigma}|\gamma') = \frac{1}{\sigma_1 \ldots \sigma_d} \mathcal{N}_{2d}(\vec{\mu}_g, \Sigma_g).$$

In the case of the subject specific posterior we assume the factorisation between subjects and parameter dependence within subject, hence we express each factor as a multivariate normal distribution

$$Q(\vec{x}_n|\gamma''_n) = \mathcal{N}_d(\vec{\mu}^n_x, \Sigma^n_x)$$

In summary, we are performing an approximate inference based on approximate posterior by maxmising evidence lower bound with respect to prior parameters $\eta$ and posterior parameters $\gamma$.

[8]:
```
infer_ch = Inferrer(agent, stimulus, responses, mask=mask,␣
 ↪fixed_params=fixed_params)
infer_ch.infer_posterior(iter_steps=200, parametrisation='cent-horseshoe')
```

```
Mean ELBO 19377.16: 100%|      | 200/200 [35:53<00:00, 10.84s/it]
```

[9]:
```
# plot posterior parameter values in comparison with true values
labels = ['beta', 'bias', 'gamma', 'kappa']
res = infer_ch.sample_from_posterior(labels, n_samples=999)

trans_pars_ch = res[0].melt(id_vars='subject', var_name='parameter')
```

```
loss_ch = infer_ch.loss

# plot convergence of stochastic ELBO estimates (log-model evidence)
plt.figure()
plt.plot(loss_ch[-150:])

fig, axes = plt.subplots(1, len(labels), figsize=(15, 5))
m = res[0].groupby('subject').mean()
std = res[0].groupby('subject').std()

for i,l in enumerate(labels):
    axes[i].errorbar(x_sample[l], m[l], std[l], c='r', marker='o', linestyle='')
    axes[i].plot(x_sample[l], x_sample[l], 'k--')
    axes[i].set_title(l)
    axes[i].set_xlabel('true value')

axes[0].set_ylabel('estimated value')
```

[9]: Text(0, 0.5, 'estimated value')

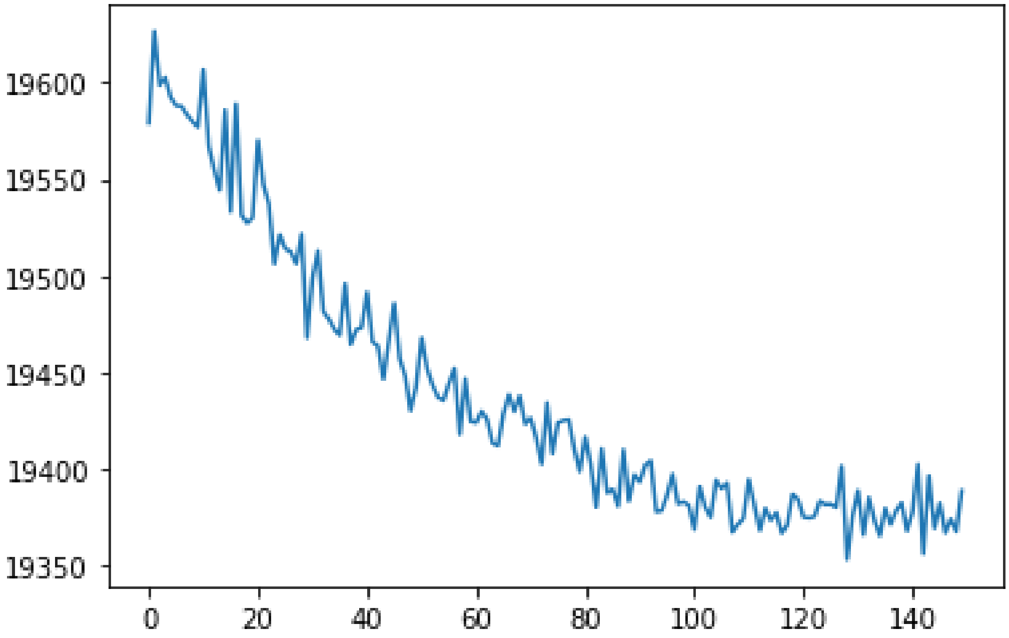

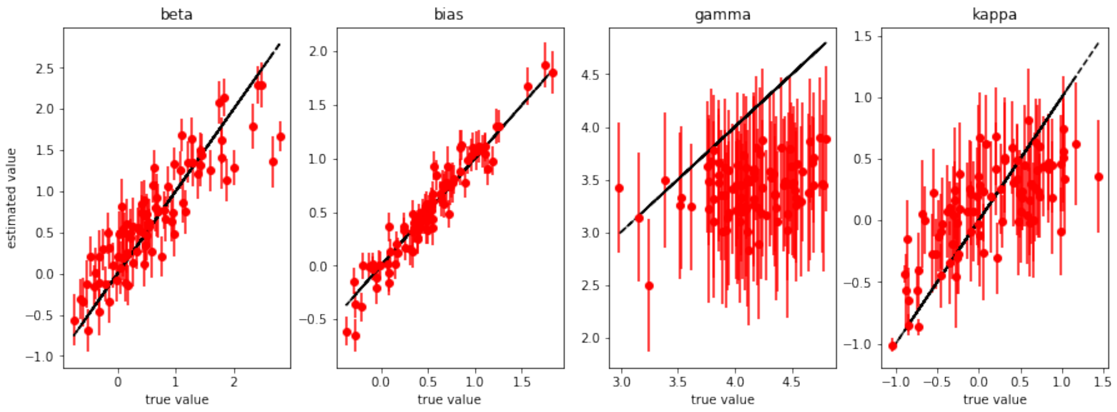


[10]:
```
names = ['beta', 'bias', 'gamma', 'kappa']
true_vals = x_sample.values.T
posterior_accuracy(names, trans_pars_ch, true_vals)

fig, axes = plt.subplots(1, 4, figsize=(15, 5))
for i, l in enumerate(labels):
    R = res[0][l].values.reshape(-1, runs) < true_vals[i]
    axes[i].hist(R.sum(0), bins=20)
```

```
beta 0.8651685393258427
bias 0.9662921348314607
gamma 0.9662921348314607
kappa 0.9101123595505618
```

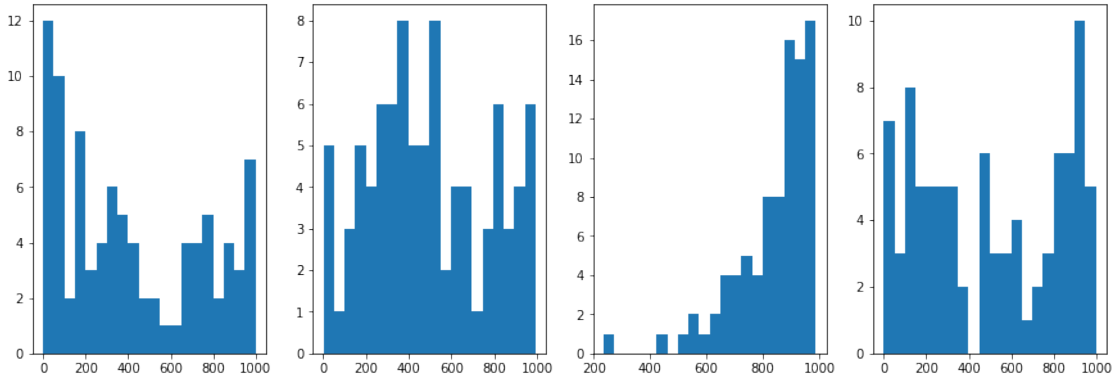

Finally we will test the horseshoe+ prior which in addition to the global hyper prior over model parameters contains local (subject specific) hyper priors over prior uncertainty

$$\mu_i \sim \mathcal{N}(m_i, s_i)$$

$$\sigma_i \sim C^+(0,1)$$
$$\rho_i^n \sim C^+(0,1)$$
$$x_i^n \sim \mathcal{N}(\mu_i, \lambda * \sigma_i * \gamma^n * \rho_i^n)$$

where we introduced another set of subject specific parameters $\gamma^n$ of the hierarchical prior. Hence, $\eta = (m_1, \ldots, m_d, s_1, \ldots, s_d, \gamma^1, \ldots, \gamma^N)$.

Similarly, as for the horseshoe prior we will use here the approximate posterior that factorises on joint distribution over hyperparameters, and joint distribution over subject specific parameters. Hence,

$$Q(Z|\gamma) = Q(\vec{\mu}, \vec{\sigma}|\gamma') \prod_n Q(\vec{x}_n, \vec{\rho}_n|\gamma''_n)$$

where we use the same functional form as the above for the posterior over hyperparameters

$$Q(\vec{\mu}, \vec{\sigma}|\gamma') = \frac{1}{\sigma_1 \ldots \sigma_d} \mathcal{N}_{2d}(\vec{\mu}_g, \Sigma_g),$$

and the same functional form for the subject specific posterior

$$p(\vec{\mu}, \vec{\sigma}, \vec{x}^1, \ldots, \vec{x}^N|D) \approx \frac{1}{\sigma_1 \ldots \sigma_d} Q(\vec{\mu}, \ln \vec{\sigma}) \prod_{n=1}^N \mathcal{N}_d(\vec{\mu}_x^n, \Sigma_x^n)$$

in the case of the horseshoe prior, and as

$$p(\vec{x}_n, \vec{\rho}_n|\gamma''_n) = \frac{1}{\rho_1^n \ldots \rho_d^n} \mathcal{N}_{2d}(\vec{\mu}_w^n, \Sigma_w^n)$$

Note that in both cases we used centered parameterisation of the parametric model, as non-centered parametrisation leads to a numerical instabilities when computing gradients over free model parameters.

[11]:
```
infer_chp = Inferrer(agent, stimulus, responses, mask=mask,␣
 ↪fixed_params=fixed_params)
infer_chp.infer_posterior(iter_steps=200, parametrisation='cent-horseshoe+')
```

```
Mean ELBO 20175.06: 100%|      | 200/200 [36:01<00:00, 10.65s/it]
```

[12]:
```
# plot posterior parameter values in comparison with true values
labels = ['beta', 'bias', 'gamma', 'kappa']
res = infer_chp.sample_from_posterior(labels, n_samples=999)

trans_pars_chp = res[0].melt(id_vars='subject', var_name='parameter')
loss_chp = infer_chp.loss

# plot convergence of stochastic ELBO estimates (log-model evidence)
plt.figure()
```

```
plt.plot(loss_chp[-150:])

fig, axes = plt.subplots(1, len(labels), figsize=(15, 5))
m = res[0].groupby('subject').mean()
std = res[0].groupby('subject').std()

for i,l in enumerate(labels):
    axes[i].errorbar(x_sample[l], m[l], std[l], c='r', marker='o', linestyle='')
    axes[i].plot(x_sample[l], x_sample[l], 'k--')
    axes[i].set_title(l)
    axes[i].set_xlabel('true value')

axes[0].set_ylabel('estimated value')
```

[12]: Text(0, 0.5, 'estimated value')

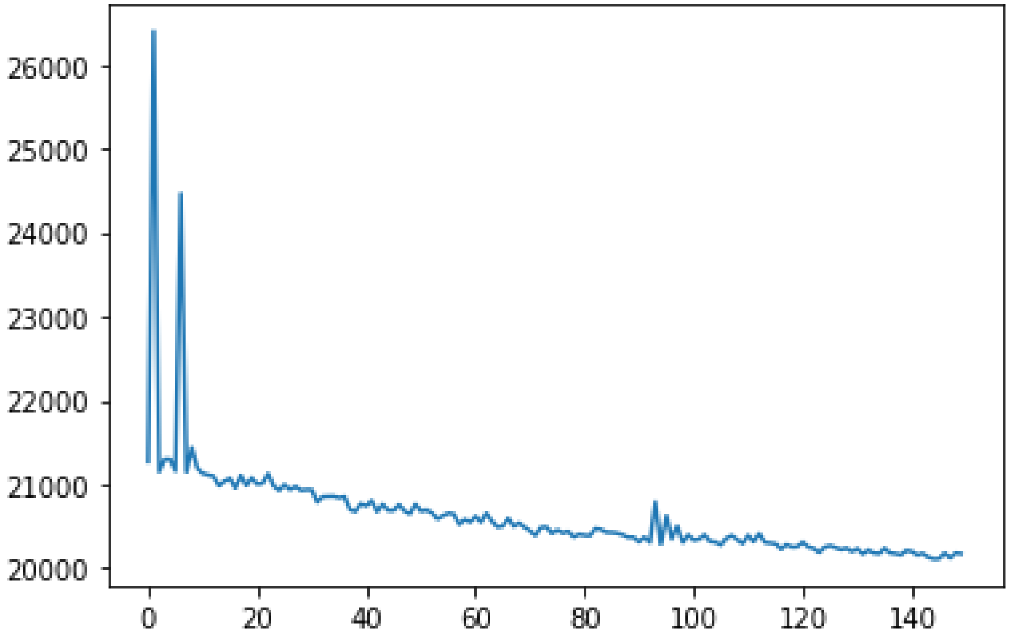

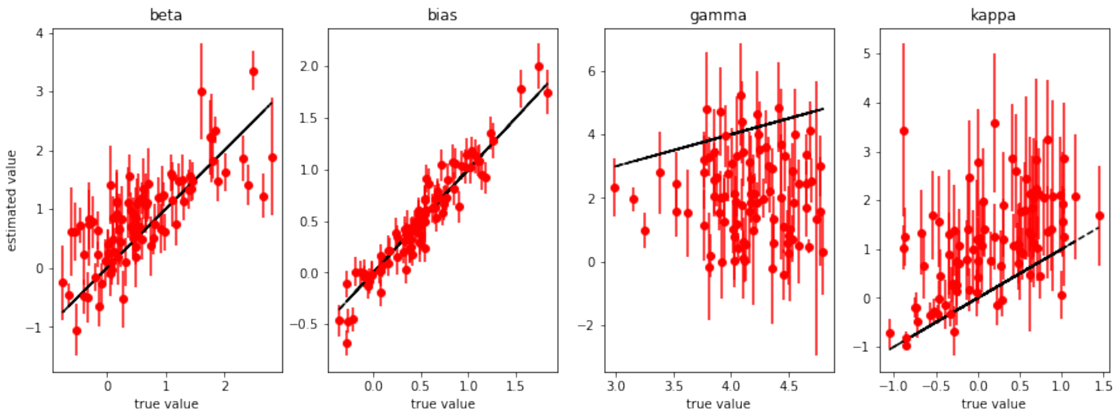


[13]:
```
names = ['beta', 'bias', 'gamma', 'kappa']
true_vals = x_sample.values.T
posterior_accuracy(names, trans_pars_chp, true_vals)

fig, axes = plt.subplots(1, 4, figsize=(15, 5))
for i, l in enumerate(labels):
    R = res[0][l].values.reshape(-1, runs) < true_vals[i]
    axes[i].hist(R.sum(0), bins=20)
```

```
beta 0.8426966292134831
bias 0.9101123595505618
gamma 0.4606741573033708
kappa 0.8651685393258427
```

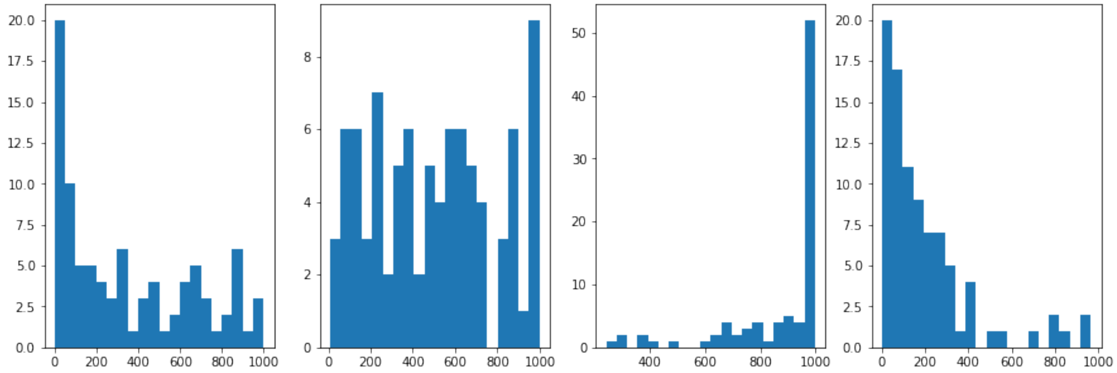
